# The crystal and cryo-EM structures of PLCγ2 reveal dynamic inter-domain recognitions in autoinhibition

**DOI:** 10.1101/2023.09.07.556539

**Authors:** Young-Cheul Shin, Ashlee Marie Plummer-Medeiros, Alison Mungenast, Hyeong-wook Choi, Karen TenDyke, Xiaojie Zhu, Jennifer Shepard, Ningning Zhuang, Liang Hu, Dongming Qian, Kangkang Song, Chen Xu, John Wang, Suresh B Poda, Maofu Liao, Yu Chen

## Abstract

Phospholipase C gamma 2 (PLCγ2) plays important roles in cell signaling downstream of various membrane receptors. PLCγ2 contains a multi-domain inhibitory region critical for its regulation, while it has remained unclear how these domains contribute to PLCγ2 activity modulation. Here we determined three structures of human PLCγ2 in autoinhibited states, which reveal dynamic interactions at the autoinhibition interface, involving the conformational flexibility of the SH3 domain in the inhibitory region, and its previously unknown interaction with a C-terminal helical domain in the core region. We also determined a structure of PLCγ2 bound to the kinase domain of fibroblast growth factor receptor 1 (FGFR1), which demonstrates the recognition of FGFR1 by the nSH2 domain in the inhibitory region of PLCγ2. Our results provide new structural insights into PLCγ2 regulation that will facilitate future mechanistic studies to understand the entire activation process.

## Introduction

Phospholipase C gamma 2 (PLCγ2, encoded by gene *PLCG2*) is a membrane-associated enzyme that catalyzes the conversion of 1-phosphatidyl-1D-myo-inositol 4,5-bisphosphate (PIP_2_) to 1D-myo-inositol 1,4,5-trisphosphate (IP3) and diacylglycerol (DAG) using calcium as a cofactor. It plays a central role in cellular signaling transduction by generating the key second messenger molecules IP3 and DAG in response to activation by a variety of transmembrane receptors or membrane-associated kinases^1,2^.

Gain-of-function variants in PLCγ2 can cause PLCγ2-associated antibody deficiency and immune dysregulation (PLAID) and autoinflammation and PLCγ2-associated antibody deficiency and immune dysregulation (APLAID) syndromes^1–7^. Mouse models of autoimmunity and autoinflammation have also been found carrying gain-of-function missense mutations in PLCγ2^8,9^. PLCγ2 is a substrate of the Bruton agammaglobulinemia tyrosine kinase (BTK) and mediates its downstream signaling. Activating mutations of PLCγ2 arise in patients with chronic lymphocytic leukemia (CLL) resistant to the BTK inhibitor ibrutinib^10,11^. Genome-wide association studies also identified a rare variant of PLCγ2 (P522R) associated with reduced risk of Alzheimer’s disease, as well as other dementias including dementia with Lewy bodies and frontotemporal dementia^12–20^. Longitudinal analysis also suggested MCI (mild cognitive impairment) patients carrying the P522R variant shows a lower rate of cognitive decline in comparison with non-carriers^21^. Activity studies indicated that this variant modestly increases the enzymatic function of PLCγ2^22–24^. Recent studies also suggest that a different missense variant of PLCγ2 (M28L) is associated with loss-of-function and an elevated risk for Alzheimer’s disease^18,25–27^.

These observations underscore the importance of finely-tuned PLCγ2 regulation. PLCγ enzymes are unique in the PLC family for containing multiple folded domains in the inhibitory region. The previously reported crystal structure of rat PLCγ1 provides insights into how the inhibitory region associates with the core region and occlude the access of the catalytic site to substrate^28^. It will be helpful to observe similar atomic details for human PLCγ2, and further address whether additional interactions may be involved in this autoinhibitory interaction.

Similar to PLCγ1^29,30^, PLCγ2 is regulated by receptor tyrosine kinases (RTKs) and transmembrane receptor-associated kinases by phosphorylation at key regulating tyrosine residues^1,2^. One of the most comprehensive studies of this regulation for PLCγ enzymes pertains to growth factor receptor tyrosine kinases phosphorylation^31–33^. Fibroblast growth factor receptor 1 (FGFR1) phosphorylates PLCγ enzymes, which contribute to the alleviation of PLCγ autoinhibition and recruitment of PLCγ to the vicinity of the membrane where its substrate PIP_2_ resides^34,35^. However, limited knowledge is available to help understand the recognition of the intact PLCγ enzyme by regulatory kinases, and if these associations alone are sufficient to trigger the major conformational changes leading to PLCγ activation.

In this study, we determined crystal and cryo-EM structures of human PLCγ2 in autoinhibited conformations (referred to as PLCγ2-C, PLCγ2-F, and PLCγ2-H), which revealed multiple recognition sites between domains in the inhibitory region and the core region. We also conducted cryo-EM analysis of PLCγ2 in complex with phosphorylated FGFR1 kinase, which confirmed that the FGFR1 kinase domain recognizes the nSH2 domain of PLCγ2 in an autoinhibition state. Our data provides a view of dynamic PLCγ2 autoinhibition, which is likely important for PLCγ2 fine regulation and applicable to PLCγ1 as well.

## Results

### Crystal structure of human PLCγ2

Multiple constructs containing N-terminal, C-terminal, or internal deletions were explored to enable protein crystallization. One construct, PLCγ2-C (14-221-GSG-239-1190, Fig. 1A), resulted in diffracting crystals. This construct contains all known structural domains of PLCγ2 (Fig. 1A, amino acid 14-1190), and a glycine-serine-glycine (GSG) peptide replacing a loop (amino acid 222-238) in the predicted EF- Hand domain.

**Figure 1.**
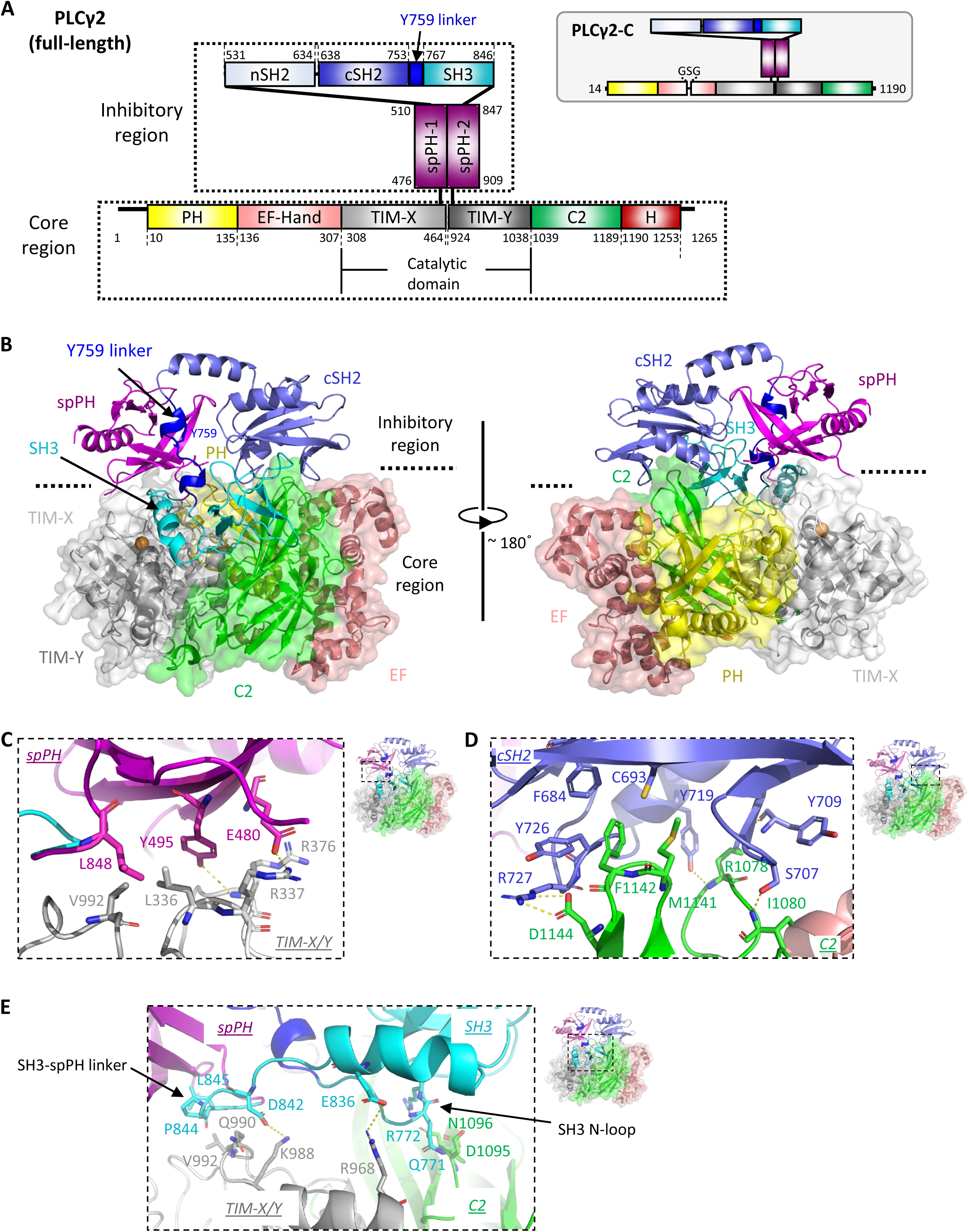
Crystal structure of human PLCγ2. **A.** Domain architecture of human PLCγ2. Color coding: PH domain – yellow; EF hand – salmon; TIM X/Y – gray; C2 domain – green; spPH domain – magenta; nSH2 domain – light blue; cSH2 domain – slate; SH3 domain – cyan; Y759 linker – bright blue. Newly observed C-terminal helices are noted as H domain and colored in red. Domain boundaries are labeled by amino acid numbers. Construct design of PLCγ2-C is illustrated in the gray box. **B.** Crystal structure of human PLCγ2 determined at 2.55 Å resolution (PLCγ2- C), presented in both front (left) and back (right) views. Domains are colored as in Figure 1A. Inhibitory region is shown in cartoon representation. The cartoon diagram of the core region is shown embedded in a surface representation. The orange sphere indicates the active site calcium cofactor. **C-E.** Interface interactions between the regulatory and core domains of PLCγ2. Domains are colored as in Figure 1A. Residues contributing to the interactions are shown in stick representation. Polar interactions are denoted by dashed lines in yellow. Regions of representation are illustrated by the dashed boxes on the complete structure on right, respectively.

A crystal structure of human PLCγ2 using this construct was determined at 2.55 Å resolution (Fig. 1B, Table S1). This particular crystal was soaked with a compound overnight before diffraction data collection. However, since no extra densities of compound were observed and the structure is identical to another one obtained without compound soaking at 2.75 Å resolution, we determined this structure at 2.55 Å resolution represents an apo structure. As it exhibited improved resolution over the structure determined at 2.75 Å resolution, we report and focus on description of this structure in this section.

The atomic model contains all domains except for the nSH2 domain, for which very limited density was observed. In this structure, the membrane engagement side of the core region is blocked by the regulatory region, indicating that it represents the autoinhibited state. The sequence connecting the cSH2 and SH3 domain (amino acid 754 – 766, Y759 linker) is clearly visible in this structure (Supplementary Fig. 1A), running toward the interface between the inhibitory and core region and interacting with the spPH domain (Supplementary Fig. 1B). This sequence contains a key regulatory tyrosine Tyr759, and the phosphorylation of Tyr759 has been shown to closely related with the activation of PLCγ2^36–38^. To mimic the potential impact of Tyr759 phosphorylation alone, we generated a phosphor-mimetic version of PLCγ2 by replacing Tyr759 with a glutamate residue. Recombinant protein of PLCγ2 Y759E exhibited enhanced activity in comparison with the wild-type PLCγ2 in a biochemical lipase assay^39^ (Supplementary Fig. 1C). These results support that phosphorylation of Tyr759 alone would be important for PLCγ2 activation.

Similar to the previously reported autoinhibited structure of rat PLCγ1 (rPLCγ1, PDB:6PBC^28^), spPH and cSH2 domains in this structure of PLCγ2 directly interact with the TIM barrel (TIM-X/Y) and C2 domain, respectively, forming major interfaces between the inhibitory and the core regions (Fig. 1C, 1D). In addition, the SH3 domain is in proximity to and interacts with both the TIM barrel and C2 domain in the core region, via the linker between the SH3 and spPH domains (SH3-spPH linker) and the SH3 N-terminal loop that follows the Y759 linker (SH3 N-loop), respectively (Fig. 1E). These interactions contribute to the interfaces between the inhibitory and the core regions and may play a role in the autoinhibition regulation as well. A direct association of the SH3 domain with the core region was not observed in the previously reported rPLCγ1 structure, in which the SH3 domain was orientated away from the inhibitory-core region interface and located on the outmost side of the inhibitory region (Supplementary Fig. 2). This may be due to the replacement of the 25-residue loop connecting cSH2 and SH3 domains by a short linker of Ser-Gly-Ser in the rPLCγ1 structure described^28^. The interface analysis between the inhibitory and core regions of human PLCγ2 suggests that three domains from the inhibitory region, spPH, cSH2, and SH3, are directly involved in the interface interaction, and probably engaged with PLCγ2 autoinhibition regulation.

### Disease-associated mutations and variants of PLCγ2

Disease-associated mutations and variants of PLCγ2 have been noted in both inhibitory and core regions, while the majority have been reported in the inhibitory region and most of them are gain-of-function mutations (Fig. 2A). When mapping the sites of gain-of-function point mutations in the autoinhibited conformation shown by the crystal structure of PLCγ2 (Fig. 2B), the majority are localized at the interface between the inhibitory and core regions, including both the cSH2/C2 and SH3-spPH linker/TIM barrel interaction sites. This observation suggests that both interaction sites are critical for PLCγ2 autoinhibitory regulation, and that mutations at either site may destabilize the inhibitory-core region interface and result in increased basal enzymatic activity. Similarly, the gain-of-function mutations harbored by two autoinflammatory disease mouse models, D993G in *Ali5* mice and Y495C in *Ali14* mice, are also located at the interface between the inhibitory and core regions, specifically the SH3-spPH linker/TIM interaction site (Supplementary Fig. 3, Supplementary Fig. 4A). We conducted IP One assays to examine the enzymatic activity of selected mutations and confirmed that they significantly enhanced basal PLCγ2 phospholipase activity (Supplementary Fig. 4B).

**Figure 2.**
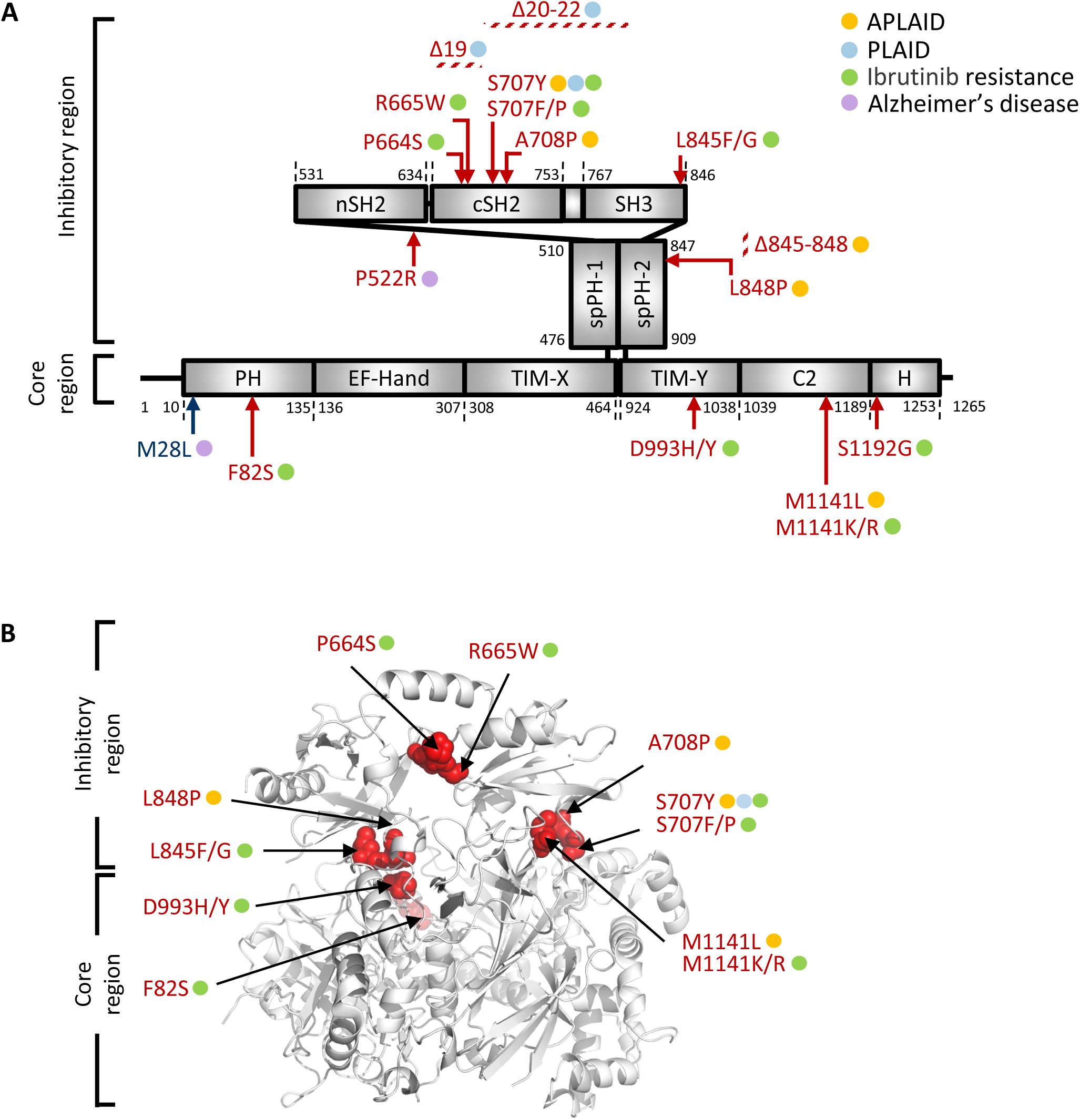
Positions and mapping of PLCγ2 mutations and variants. **A.** Positions of gain-of-function point mutations are indicated by red arrows, and the deletions are denoted by red diagonal stripes. Loss-of-function mutation is noted by a dark blue arrow. Disease association is illustrated by filled circles: yellow - APLAID; turquoise blue – PLAID; green – Ibrutinib resistance; purple – Alzheimer’s disease. **B.** Mapping of PLCγ2 gain-of-function mutations onto the crystal structure of PLCγ2. Residues at the mutation site are colored in red and shown in sphere representation to label the position of mutation. Label denotes mutations discovered so far at each position. Disease association is illustrated by filled circles as in Figure 2A. Sites for P552R and S1192G are not shown as residues Pro522 and Ser1192 are invisible or absent in the crystal structure, respectively.

When mapping the reported deletions in the crystal structure of PLCγ2, the deletion linked to APLAID (Δ845-848) is located within the SH3-spPH linker in direct association with the TIM-barrel catalytic domain in the core region (Supplementary Fig. 4C, Fig. 1E). This deletion has been shown to greatly enhance PLCγ2 activity^5^, likely due to the disruption of autoinhibitory recognition at this site. As expected, the causative deletions for PLAID (exon 19 and exon 20-22 deletions: Δ19 and Δ20-22) result in the loss of the most of cSH2 domain (amino acid 646-685) and the majority of cSH2-Y759 linker-SH3 domains (amino acid 686-806), respectively, which are expected to have a major impact on the integrity of autoinhibitory interface and result in higher basal phospholipase activity (Supplementary Fig. 4C).

Our analysis suggests that the inhibitory-core region interface has crucial functions in PLCγ2 autoinhibition regulation and activity modulation. The recognition sites between the cSH2 and C2 domains, and between SH3-spPH and TIM barrel, respectively, play indispensable roles.

### Structure analysis by single-particle cryogenic electron microscopy (cryo-EM)

To capture additional structure details in full-length PLCγ2 and gain insights into PLCγ2 recognition by FGFR1, we generated recombinant proteins of human PLCγ2 with the complete sequence (Fig.1A, amino acid 1-1265) and the kinase domain (amino acid 456-774) of human FGFR1 (FGFR1K). The association between PLCγ2 and FGFR1K was examined by size exclusion chromatography in the presence and absence of FGFR1K phosphorylation of the key regulating tyrosine Tyr766. Both full-length PLCγ2 and PLCγ2-C were tested. The results demonstrated that phosphorylated FGFR1K (pFGFR1K) interacts and co-elutes with PLCγ2 in both cases, while FGFR1K lacking phosphorylation is largely eluted separately from PLCγ2, confirming that phosphorylation of FGFR1K is required for associating with PLCγ2 (Supplementary Fig. 5) as previously reported^34,35^.

For cryo-EM analysis, the full-length PLCγ2 was incubated with pFGFR1K. A 3D reconstruction of PLCγ2 was obtained at 3.7 Å resolution (referred to as structure PLCγ2-F, Supplementary Fig. 6), but the SH3 and C-terminal regions of PLCγ2 and FGFR1K were not well resolved. To improve the density, the particle set was further processed in two directions (Supplementary Fig. 6). Firstly, we performed masked 3D classification without alignment in Relion and local refinement in cryoSPARC, obtaining a 4.1 Å resolution map of PLCγ2 with clear density of the C-terminal region (referred to as structure PLCγ2-H). Secondly, we generated a complex mask covering both the densities of PLCγ2 and FGFR1K, obtaining a full complex map at 4.0 Å resolution (referred to as structure PLCγ2/pFGFR1K) by performing 3D classification without alignment and local refinement (in Relion and in cryoSPARC, respectively). Three cryo-EM structures involving human PLCγ2 were determined (Supplementary Fig. 6, Table S2).

### Cryo-EM structures of PLCγ2

All major domains of PLCγ2 were mostly well-resolved in the cryo-EM density map of PLCγ2-F, including the nSH2 domain in the inhibitory region that was missing in the crystal structure (Fig. 3A). The nSH2 domain is connected to the cSH2 domain and positioned to readily bind to upstream kinases and recruit interacting partners, as previously observed in the crystal structure of rPLCγ1^28^. Overlay of the cryo-EM structure of PLCγ2 with the crystal structure of rPLCγ1 revealed a ∼ 20֯ shift of the nSH2 domain (Supplementary Fig. 7A), which may be related to its conformational flexibility relative to the rest of the protein.

**Figure 3.**
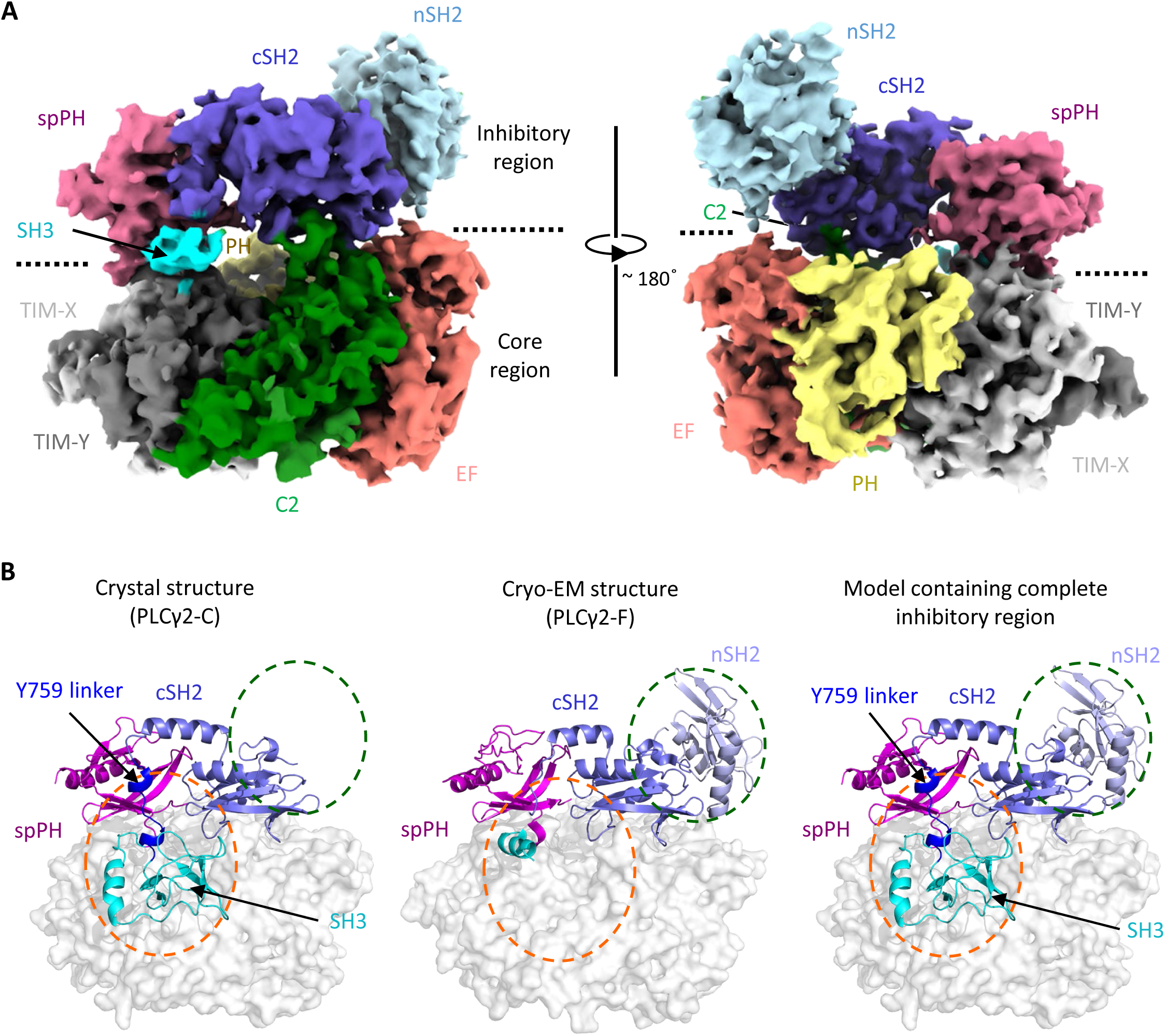
Cryo-EM structure of full-length human PLCγ2 (PLCγ2-F) **A.** Cryo-EM map of PLCγ2-F (3.7 Å resolution) presented in both front (left) and back (right) views. Domains are colored as in Figure 1A, and the nSH2 domain (in light blue) is visible. **B.** Structural model of PLCγ2 with complete inhibitory region. Left: crystal structure of PLCγ2 contains the entire SH3 domain and Y759 linker (highlighted in orange circle) but not the nSH2 domain. Middle: cryo-EM structure of full-length PLCγ2 (PLCγ2-F) has visible nSH2 domain (highlighted in green circle) but misses the vast majority of SH3 and Y759 linker. Right: a structural model of PLCγ2 containing all inhibitory domains was built by overlaying domains visible in both structures. Core region is shown in surface representation in grey. Inhibitory region domains are shown in carton representation and colored as in Figure 1A.

The cryo-EM map of PLCγ2-F contains weak densities for the SH3 domain and the Y759 linker connecting cSH2 and SH3 domains, indicating that this region may be conformationally flexible. Domains visible in both crystal and this cryo-EM structure of PLCγ2 are well overlayed (R.M.S.D. = 1.262 Å), which allows us to build a structure model containing all inhibitory domains, providing a comprehensive view of PLCγ2 in the autoinhibited state (Fig. 3B). In this model, the membrane-associating side of the core region is engaged in direct interactions with three out of the four major domains in the inhibitory region (spPH, SH3, and cSH2), while the fourth domain, the nSH2 domain, is free from the interface engagement and accessible for recognition with interacting partners.

The cryo-EM map of PLCγ2-H contains strong densities for the SH3 domain as well as extra helical densities that were not explained by the known domains from PLCγ2 (Fig. 4A, and Supplementary Fig. 6). The extra densities connect to and follow the C-terminal end of the C2 domain in the core region, indicating that they correspond to the very C-terminal sequence of the full-length protein following the C2 domain (Fig. 1A). The helical conformation of this region was also predicted by AlphaFold (Supplementary Fig. 7B), in agreement with our analysis. An atomic model was built into the visible densities in this region, suggesting that the C-terminal sequence 1189 to 1259 forms a helix-turn-helix structure (Fig. 4A). As this region has not been structurally annotated to our knowledge, we named this novel structure the H domain (Fig. 1A and Fig. 4A). In this cryo-EM structure, the H domain folds back to the rest of the protein and contacts the SH3 domain, suggesting a role for such an interaction in regulating PLCγ2 autoinhibition and activation.

**Figure 4.**
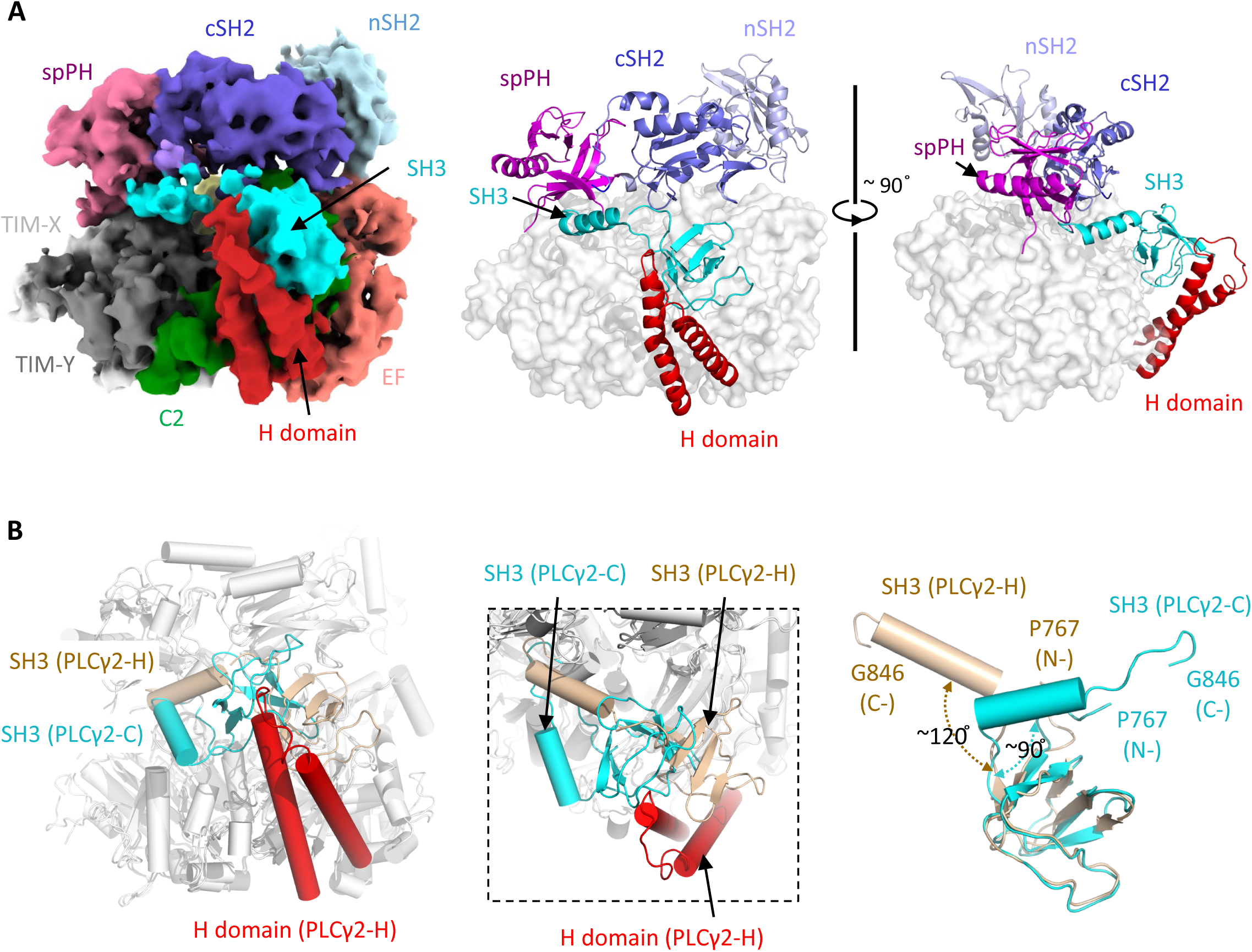
Cryo-EM structure of full-length PLCγ2 containing visible C-terminal helical structure, PLCγ2- H. **A.** Cryo-EM structure PLCγ2-H revealed the C-terminal end forms a double helical structure (newly named H domain) and makes direct contact with SH3 domain in the inhibitory region. Left: Cryo-EM map of PLCγ2-H. Domains are colored as in Figure 1A, and the C-terminal H domain is colored in red. Middle and right: front (middle) and side (right) views of PLCγ2-H structure. C-terminal helices are shown in cartoon representation in red, and the rest of the core region is shown in surface representation in grey. Inhibitory region domains are shown in cartoon representation as colored previously. **B.** Conformational flexibility of the SH3 domain in the presence of C-terminal helices. Left: overlay of crystal structure of PLCγ2 (PLCγ2-C) with cryo-EM structure PLCγ2-H highlighted conformational difference of SH3 domain in full-length protein representation. The SH3 domain is shown in cyan in PLCγ2-C, and the SH3 domain and H domain are colored in wheat and red in PLCγ2-H, respectively. The rest of the structures are shown in grey. Middle: focused representation on SH3 domains from the left overlay. Right: overlay of SH3 domain structures from PLCγ2 crystal structure PLCγ2-C (cyan) and cryo-EM structure PLCγ2-H (wheat) by its N-terminal beta-sheet region (amino acid 767-828). The orientation between N- and C-terminal region of SH3 is illustrated by the relative angel and shown by the dashed arrow. Positions of first (N-) and last (C-) residues of this region are denoted.

Superimposition of the cryo-EM structure PLCγ2-H with the crystal structure PLCγ2-C revealed conformational changes in the SH3 domain (Fig. 4B). Although the structural fold of the N-terminal region of the SH3 domain (amino acid 767-828) remains unchanged, it shifts and rotates toward the H domain in PLCγ2-H. The C-terminal helical region of the SH3 domain (amino acid 829-846) also rotates and displays a more extended conformation in PLCγ2-H. The N- and C- terminal regions of the SH3 domain are ∼ 90֯ relative to each other in PLCγ2-C, while this angle is extended to ∼ 120֯ in PLCγ2-H (Fig. 4B). This observation is consistent with the conformational flexibility of SH3 domain as noted in the structure PLCγ2-F, which may contribute to the direct association between the SH3 and H domains and be relevant to its role in PLCγ2 activity modulation.

The crystal and cryo-EM structures described so far likely represent different auto-inhibited conformations of human PLCγ2, and we decided to further examine this in the presence of interacting partners.

### Cryo-EM structure of PLCγ2 in complex with FGFR1K

The cryo-EM density map of PLCγ2/pFGFR1K was refined to 4.0 Å average resolution and confidently demonstrated the presence of FGFR1K in the complex (Fig. 5A). The recognition of FGFR1K by PLCγ2 is mainly mediated by the nSH2 domain, in a similar orientation as previously reported crystal structure of FGFR1K bound to rat PLCγ1 nSH2 (PDB: 3GQI^32^). Conformational flexibility of nSH2 domain was observed when comparing PLCγ2 structures obtained so far, which may facilitate its recognition by regulating partners (Supplementary Fig. 7A).

**Figure 5.**
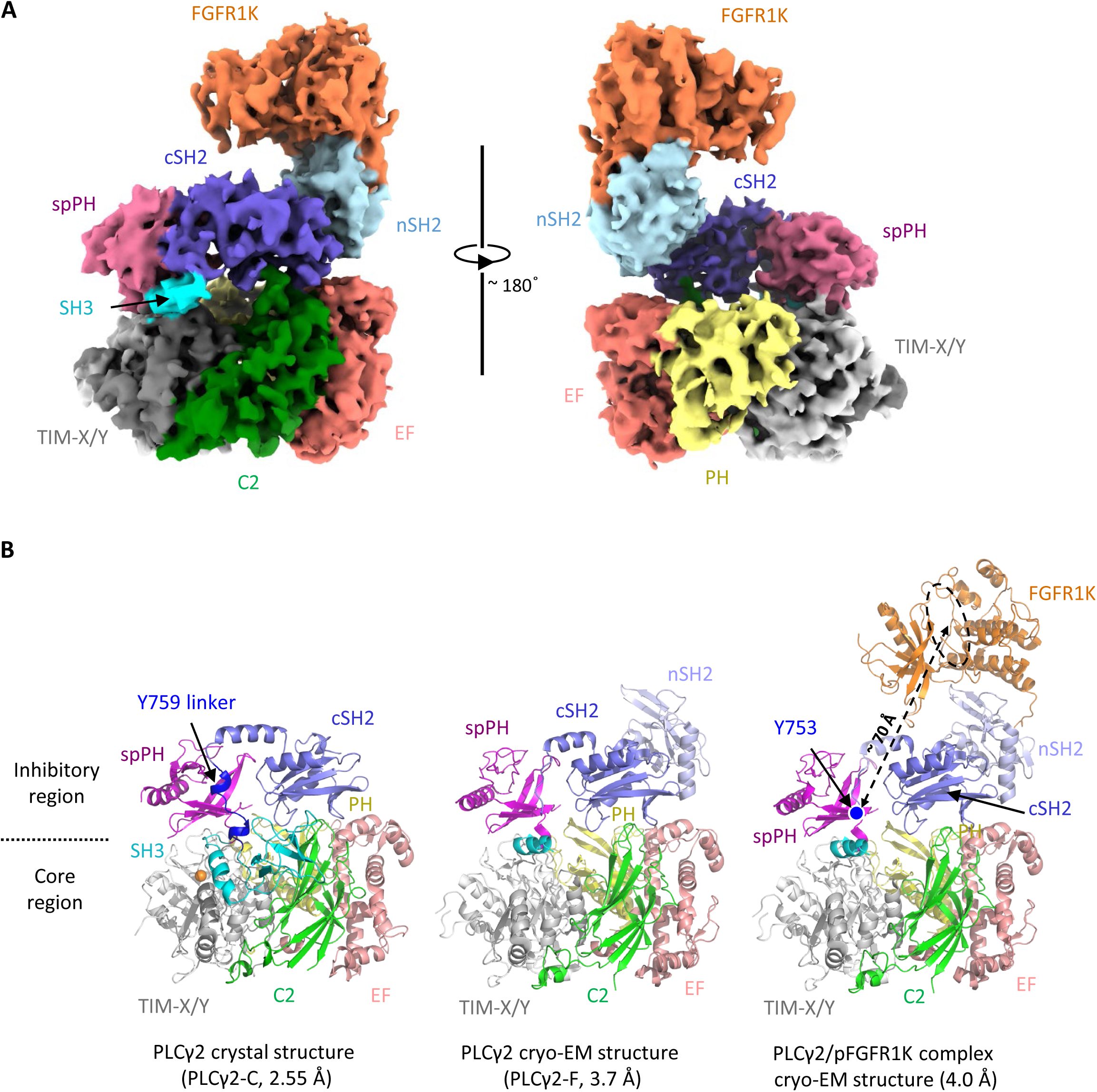
Cryo-EM structure of human PLCγ2 in complex with phosphorylated FGFR1K. **A.** Cryo-EM map of human PLCγ2/pFGFR1K complex (4.0 Å resolution) presented in both front (left) and back (right) views. PLCγ2 domains are colored as in Figure 1A, and FGFR1K is colored in orange. **B.** Structures of PLCγ2 and PLCγ2/pFGFR1K complex. All shown in cartoon representation and colored as in Figure 5A. Active site of FGFR1K is noted by a black circle in dashed line. The distance between Y753 in the cSH2-SH3 linker and the active site of FGFR1K is noted by a dashed arrow. PLCγ2 cryo-EM structure only shows PLCγ2-F for illustration.

PLCγ2 exhibited an autoinhibited conformation in this complex structure, similar to the conformation observed in the PLCγ2 alone crystal and cryo-EM structures (Fig. 5B). This suggests that the physical association of FGFR1K with PLCγ2 alone may not induce a prominent conformational change in PLCγ2. Previous cryo-EM analysis of a cross-linked PLCγ1-FGFR1K complex also showed an autoinhibited state of PLCγ1 in the complex^31^. Key regulatory residues Tyr753 and Tyr759 of PLCγ2 are expected to be phosphorylated by regulating kinases, which may contribute to subsequent autoinhibition relief and enzyme activation. Tyr759 was not visible in the cryo-EM structure of PLCγ2/pFGFR1K, making it difficult to evaluate how this residue may be accessible to the active site of FGFR1K. Tyr753 is visible in this structure and is about 70 Å in linear distance to the active site of FGFR1K (Fig. 5B). Based on this observation, Tyr759 and Tyr753 are unlikely directly accessible to the active site of FGFR1K, and additional conformational changes would be needed to allow the phosphorylation of either tyrosine by bound FGFR1. As endogenous FGFR1 is a transmembrane protein located on the plasma membrane and PLCγ2 typically executes enzymatic activity on the membrane, it may be critical to fully capture the modulation of PLCγ2 by FGFR in the presence of relevant membrane environments.

Our results suggest that phosphorylated FGFR1K recognizes the nSH2 domain in the autoinhibited PLCγ2, which may require additional conformational modulation to enable the phosphorylation of key regulating tyrosine residues and thus activate PLCγ2.

## Discussion

In comparison with other members of the phospholipase C family, the PLCγ enzymes contain a structurally complicated inhibitory region. Our results suggest that most of these domains, except for the nSH2 domain, are directly involved in the association with the core region in the autoinhibited states. The structures we determined exhibit a wide range of conformations that probably exist during autoinhibition, indicative of dynamic inter-domain interactions at the interface between the inhibitory and core regions that likely contribute to the regulation of PLCγ2 by different processes (Figure 6, left). Two recognition areas, the site involving SH3-spPH from the inhibitory region and the TIM barrel catalytic core, and the interface between the cSH2 domain in the inhibitory region and C2 domain in the core region, are present in all auto-inhibited structures, suggesting they may be required for the enzyme in the fully closed, autoinhibition state. Most of the gain-of-function mutations are also located in these areas, supporting the integrity of both recognition sites may be critical for complete inhibition of PLCγ enzymes (Figure 2B, ^28^).

**Figure 6.**
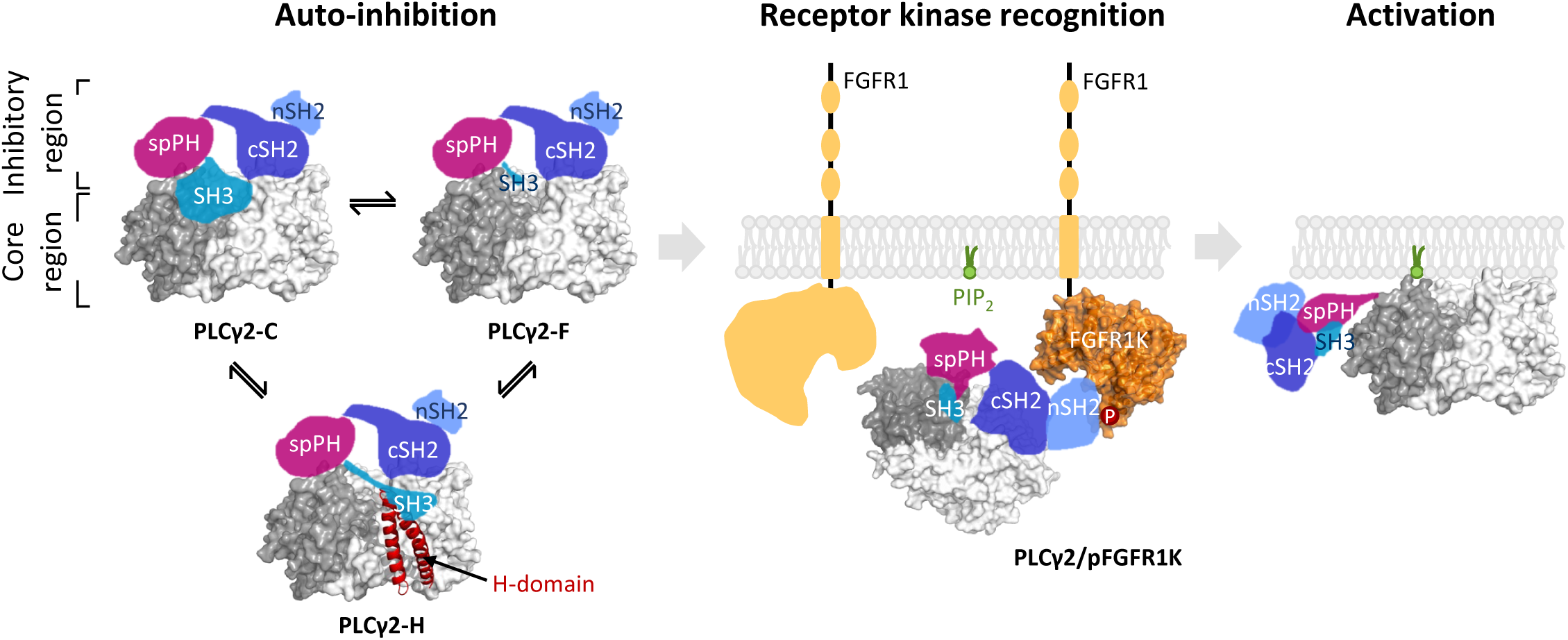
Dynamic interactions in autoinhibited states of PLCγ2 and recognition by receptor kinase FGFR1. The auto-inhibited states of PLCγ2 (left) exist in multiple conformations, exhibited by structural flexibility of the SH3 and C-terminal H domains. The recognition sites between cSH2 and C2 domains, and between SH3-spPH linker and TIM barrel, respectively, play indispensable roles. Recognition of PLCγ2 by FGFR1K (middle) is mediated by the nSH2 domain in the inhibitory region, and FGFR1K phosphorylation dependent. This recognition may position PLCγ2 in proximity to the membrane as well as a second FGFR1K nearby, both of which may be involved in the next step of phosphorylation and conformation change of PLCγ2, resulting in the disengagement of the inhibitory region and access of catalytic core to PIP_2_ substrate the membrane (right). Core region is shown in surface representation in grey, in which the TIM barrel that harbors the catalytic center is colored in dark grey. Newly identified H domain is shown in cartoon representation in red. Inhibitory region domains are illustrated in color blocks and colored as in Figure 1A. FGFR1K associated with PLCγ2 is presented in surface representation in orange, and the rest of the molecule and a second copy of FGFR1 are shown in orange shape blocks. The phosphorylated site of FGFR1K is denoted by “P” in red circle. Plasma membrane bilayer is shown in light grey, and a substrate PIP_2_ is highlighted in green.

Both the SH3 domain in the inhibitory region and the newly observed H domain in the C-terminal region of the core domain contribute to the observed structural dynamics in PLCγ2 (Figure 6, left). The SH3 domain exhibited different conformations among the three PLCγ2 structures we present in this manuscript. It interacts with different domains from the core region and is covalently connected to the regulatory Y759 linker and catalytic core-covering spPH domain, indicating that the SH3 domain may play an important role in connecting different domains during the PLCγ2 autoinhibition regulation. This may be applicable to PLCγ1 as well. It remains to be investigated mechanistically how the conformational flexibility of the SH3 domain directly impacts the activity regulation of PLCγ enzymes. One structure from our cryo-EM studies, PLCγ2-H, also showed, for the first time, structural details of the C-terminal region of PLCγ2. It exhibited a helix-turn-helix structure and folds back to the rest of the core region, engaging directly in interactions with the SH3 domain from the inhibitory region. We named this segment the H domain to reflect its structural features, and the structure indicates that this newly-described domain may also participate in autoinhibition and perhaps activity regulation of PLCγ2. Interestingly, S1192G, one of the reported gain-of-function mutations arising from Ibrutinib resistance^11^, is localized in this segment. Studies are still needed to elucidate how this structure of PLCγ2 and its recognition to the inhibitory region contributes to the enzymatic activity regulation in cells. It remains unclear whether similar structures exist at the C-terminal region of PLCγ1. This segment has moderate sequence similarity between human PLCγ1 and PLCγ2 (∼17% sequence identity, Supplementary Fig. 3), and *in silico* predictions suggests a high likelihood of disorder in this segment of PLCγ1.

PLCγ enzymes can be activated downstream of activated FGFR signaling, while the mechanistic understanding of this process is still missing. The cryo-EM structure of PLCγ2 in complex with the phosphorylated kinase domain of FGFR1 shows that their recognition is mediated by the nSH2 domain in the inhibitory region, consistent with similar reports for PLCγ1^31,32^. This association has a very limited impact on the overall conformation of PLCγ2, suggesting this may represent the first step of recognition (Figure 6, middle). This will also likely bring PLCγ2 in close proximity to the inner leaflet of the plasma membrane, where its substrate PIP_2_ resides.

It also remains unclear how FGFR1K phosphorylates the key regulating tyrosine residues in PLCγ enzymes. Our cryo-EM structure of PLCγ2/pFGFR1K suggests these residues remain far from the active site of the associating FGFR1K (Figure 5B). As FGFR1R dimerizes upon activation, there may be a possibility for the involvement of the kinase domain from the other copy of FGFR1 in getting access to and modifying these key tyrosine residues located in the Y759 linker between the cSH2 and SH3 domains in the inhibitory region (Figure 6, middle). Upon phosphorylation, this linker may disengage from its interaction with the spPH domain and induce further conformational changes to remove the inhibitory region from the membrane-associating side of the core region (Figure 6, right). In this case, the proximity of PLCγ2 to the plasma membrane as a result of the FGFR1K- PLCγ2 interaction may additionally facilitate the direct association of the membrane with the expected lipid-associating domains in the core region (PH, EF-Hand, and C2 domains), which will further enhance the anchoring of PLCγ2 to the membrane and its access to the PIP_2_ substrate.

Our studies provide novel structural insights into PLCγ2 autoinhibition and upstream factor recognition, which may have general mechanistic implications for PLCγ enzymes in healthy and disease states.

## Method and materials

### Protein cloning, expression, and purification

The human PLCγ2 construct PLCγ2-C (14-221-GSG-239-1190) was generated to facilitate protein crystallization, and cloned into a modified pFastBacHT vector, pFBLIC, which incorporates a His6-tag at the amino terminus of the expressed protein cleavable by TEV protease, using a previously published ligation-independent cloning (LIC) strategy^40^. The construct was transformed into DH10Bac cells (Invitrogen) to produce bacmid DNA, which was subsequently used to generate baculovirus in Sf9 cells according to the manufacturer’s protocol (Invitrogen Bac-to-Bac manual). High Five insect cells at a density of 2.0 X 10^6^ cells/ml were infected with baculovirus encoding PLCγ2-C at an MOI of ∼5.0 for 48 hours at 27 °C. Cells were harvested at 2000 rpm and resuspended in lysis buffer containing 25 mM Tris pH 7.5, 300 mM NaCl, 5% glycerol, 1 mM TCEP and supplemented with EDTA-free complete protease inhibitor tablets (Roche Applied Science). Cells were sonicated and then spun at 12000 rpm for 1h at 4 °C. The clarified supernatant was loaded onto a Ni-NTA column (Qiagen) that equilibrated with lysis buffer. The column was washed with 20 column volumes of the buffer containing 20 mM imidazole and then eluted with 20 column volumes of the buffer containing 250 mM imidazole. Eluted PLCγ2 protein was dialyzed overnight against lysis buffer in the presence of the TEV protease to remove the His6 tag. Cleaved protein was then subjected to a second passage over a Ni-NTA column. Fractions containing purified proteins were concentrated and further purified by a Superdex 200 size-exclusion column (GE Healthcare) equilibrated in buffer containing in the 25 mM Tris pH 7.5, 150 mM NaCl, 1 mM TCEP, and 5% glycerol.

For cryo-EM and biochemical studies, the DNA sequence encoding full-length human PLCγ2 (1-1265) was cloned into the pTT5 expression vector with N-terminal Flag tag to facilitate protein purification. The construct was transiently transfected into Expi293F^TM^ cells (Thermofisher). The cells were cultured at 37 °C in Reduced-Serum Medium (Gibco) with 8% CO_2_ at a speed of 125 rpm, and cells were collected 72 hours after transfection. The cells were harvested at 2000 rpm and resuspended in lysis buffer containing 25 mM Tris pH7.5, 300 mM NaCl, 5% glycerol, and supplemented with EDTA-free complete protease inhibitor tablets (Roche Applied Science). Cells were sonicated and then spun at 12000 rpm for 1 hour at 4 °C. The clarified supernatant was captured by Anti-Flag M2 affinity gel resin (Sigma), followed by size-exclusion chromatography purification using a Superdex 200 column (GE Healthcare) equilibrated in buffer containing 25 mM Tris pH 7.5, 150 mM NaCl, 1 mM TCEP, and 5% glycerol. The recombinant protein of human PLCγ2 Y759E was generated using the same procedure.

To generate FGFR1 kinase domain recombinantly, plasmid DNA encoding the kinase domain (residues 456-774) of human FGFR1 with six substitutions (Y463F, Y583F, Y585F, Y653F, Y654F, Y730F)^41^ was cloned into pET28a vector incorporating a His6-tag and a SUMO tag at the amino terminus of the expressed protein and expressed in *E.coli* BL21 gold strain (Novagen). Cells were grown at 37°C in TB medium containing 0.1 mg/ml kanamycin and 0.050 mg/ml tetracyline to an OD600 of ∼3.0. Protein expression was induced for 20 hours at 16 °C with 0.1 mM IPTG (final concentration). Cells were collected by centrifugation and resuspended in lysis buffer containing 25 mM Tris pH 7.5, 300 mM NaCl, 5% glycerol, and 1 mM TCEP. The cells were lysed using high pressure homogenizer and CHAPS was added to a final concentration of 0.5% w/v followed by incubation at 4 °C for 30 min. Soluble lysate was prepared by centrifugation and the kinase domain of FGFR1 was isolated by Ni-NTA column (Qiagen).

The eluted protein was dialyzed overnight against lysis buffer in the presence of the ULP1 protease to remove the His-SUMO tag, and then subjected to a second passage over a Ni-NTA column. The unbound protein was further purified by ion-exchange chromatography using a HiTrap^TM^ Q-HP 5 mL column (Cytiva), and the phosphorylated kinase domain of FGFR1 was collected after size exclusion chromatography purification using Superdex 75 10/300 GL column (GE Healthcare). To prepare the dephosphorylated form, purified phosphorylated kinase domain of FGFR1 was treated with His-tagged lambda phosphatase in the presence of 2 mM MnCl_2_ at 4 °C overnight, followed by isolation using Ni-NTA column and purification by size exclusion chromatography (Superdex 75 column). Intact protein molecular weight was detected by LC-MS using an Agilent 6530 Q-TOF mass spectrometry connected to an Agilent 1290 UPLC, confirming the phosphorylation status. The phosphorylation site in the phosphorylated kinase domain of FGFR1K was detected at Tyr766 by Protein Mapping.

Protein concentrations were determined using the A280 method with extinction coefficients calculated using the ProtParam tool (ExPaSy Molecular Biology Server^42^). The prepared protein was further analyzed by SDS-PAGE to confirm concentration and purity, and subsequently stored at -80 °C after flash freezing in liquid nitrogen.

### Crystallization

Crystals of PLCγ2 were grown initially by sitting drop vapor diffusion using purified protein from construct PLCγ2-C. Protein solution was prepared at 10 mg/ml in buffer containing 20 mM HEPES pH7.5, 150 mM NaCl, 1 mM TCEP, and 5% glycerol. Two hundred nanoliters of this protein solution was mixed with 200 nanoliters of reservoir solution that contains 0.1 M magnesium formate and 15 % w/v PEG 3350 and equilibrated against a 40 µl reservoir. Crystals appeared after one week at 18 °C. Diffraction quality crystals were grown at 18 °C by microseeding hanging drops. Drops were prepared by mixing 1 µl of reservoir solution (0.1 M magnesium formate, 15 % w/v PEG 3350) with 1 µl of protein solution.

Drops were equilibrated against a 500 ml reservoir. Crystals were grown to ∼100 µm on the longest edge after 4-5 days. Some of them were also transferred to a new 2 µl hanging drop with a reservoir solution (0.1 M magnesium formate, 15 % w/v PEG 3350) for overnight compound soaking. The crystals were cryo-protected in mother liquor containing 20% ethylene glycol and flash-frozen in liquid nitrogen on nylon loops.

### Diffraction data collection and crystal structure determination

The diffraction data was collected at Shanghai Synchrotron Radiation Facility (SSRF, BL19U1^43^) and processed using autoPROC^44^ that includes XDS^45^ and AIMLESS (CCP4 package)^46^. The initial molecular replacement was performed by Phaser MR^47^ with rat PLCγ1 core domain (PDB code: 6PBC) as the template. Structure refinement was carried out using Refmac5^48^ combined with several rounds of COOT^49^ manual fitting. The calcium ion and antimony ion were accurately fit into the positive density maps according to the electron density maps after an initial round of refinement. Figures were prepared with PyMOL^50^ (The PyMOL Molecular Graphics System). Complete data collection and refinement statistics are shown in Table S1.

### Cryo-EM sample preparation and data collection

The final preparation of the PLCγ2 sample was carried out using a Superose 6 Increase 10/300 column, which was equilibrated with 25 mM Tris (pH 7.9) and 150 mM NaCl. Fractions were collected, concentrated to 1 mg/ml, and then mixed with 5 mg/ml of FGFR1K at a 1:18 molar ratio. Following a 30- minute incubation, the sample was further concentrated to 1.06 and 2.7 mg/ml for cryo-EM grid preparation. A volume of 3.5 μl of the prepared sample was applied to a glow-discharged holey grid (Quantifoil Au300 R1.2/1.3), and then blotted for 3.5 seconds at room temperature with 90% humidity using a Gatan Cryoplunge 3 system. The samples were then vitrified in liquid ethane cooled by liquid nitrogen. Cryo-EM images were collected on a Talos Arctica (Thermo Fisher Scientific). Additional details are provided in Table S2.

### Cryo-EM data processing

Detailed information regarding the number of particles and map resolutions can be found in Table S2 and Supplementary Figure 6. Image sets from two different samples were combined, consisting of 3,369 images from the 1.06 mg/ml sample and 2,974 images from the 2.7 mg/ml sample. Movies were motion corrected with MotionCor2^51^, and defocus values calculated with CTFFIND4^52^. All initial two-dimensional (2D) processing steps were performed using SAMUEL (Simplified Application Managing Utilities for EM Labs) Package^53^. Dose-weighted images were binned 3x with “sampilcopy2d.py” and interactively screened using “SamViewer”. Particle picking, particle stack generation, star file generation and 2D classification were performed using “samautopick.py”, “sampilboxparticle.py”, “samrelion.py new” and “samtree2dv3.py”, respectively. Additional 2D classification from individual 2D classes was performed using “samrelion.py 2dcls” which utilized RELION 3.0, and 3D classification was carried out in RELION 3.0^54^.

For mask generation, the best maps from 3D classification were low-pass filtered to 30 Å resolution using “relion_image_handler” and masks were generated using “relion_mask create” in RELION with 2 pixels extension and 6 pixels for smoothening of the edges. Further refinement for obtaining final maps was carried out using “Local refinement” in cryoSPARC^55^. The resolutions were estimated based on the FSC=0.143 criterion, and the curves were calculated using using “Comprehensive Validation” in Phenix^56^. Finally, the processed maps were obtained for analysis, and the resulting structures were referred to as PLCγ2-F, PLCγ2-H, and PLCγ2-FGFR1K, as described in Supplementary Fig. 6.

### Cryo-EM structure model building, refinement, and display

Three structure models (PLCγ2-F, PLCγ2-H, and PLCγ2/pFGFR1K) from cryo-EM data were built using Alphafold2 models of PLCγ2 and FGFR1K as the starting templates. SH2, SH3, and C-terminal helix domains were extracted using PyMOL (Version 2.5.2, Schrödinger, LLC.), and all fragmented domains were fitted to the maps using the “fit in map” function in UCSF ChimeraX^57^. Any disconnected loops, side chain rotamers, varied secondary structures, or unmatched regions to the map were manually corrected at sigma levels of 3.0∼7.0 using “Real space refine zone” in Coot^58^. The models were further refined using “real-space refinement” in Phenix. Final models were validated using “Comprehensive validation” in Phenix after outlier correction in Coot. Figures of structural models and cryo-EM maps were generated using UCSF ChimeraX and PyMOL.

### Size exclusion chromatography examining PLCγ2/FGFR1K association

Phosphorylated and dephosphorylated FGFR1K were mixed in equal molar ratio with PLCγ2 (full-length PLCγ2 and PLCγ2-C) respectively in the presence of 5 mM MgCl_2_ and incubated at 4 °C overnight. The mixtures were then loaded onto a Superdex 200 10/300 GL size exclusion column equilibrated with buffer containing 25 mM Tris pH 7.5, 150 mM NaCl, 5 mM MgCl_2_, 5% glycerol, and 1 mM TCEP, and the elution fractions were analyzed by SDS-PAGE.

### Lipase activity measurement

Lipid vesicles containing the fluorogenic substrate XY-69 was prepared as previously described^39,59^. Briefly, lipid vesicles were generated containing 0.46 µM XY-69, 57.6 µM phosphatidylinositol 4,5- bisphosphate (PIP2), and 230 µM phosphatidylethanolamine. In each reaction, 10 µl of lipid vesicle solution was mixed with 2 µl of recombinant PLCγ2 (3.13 ng), and the fluorescence was read immediately and recorded at 2 minutes intervals for 64 minutes (485 nm excitation, 520 nm emission).

The assessment of PLC activity using the IP One assay was carried out by co-transfecting HEK293 cells (*PLCG1* knock-out) with constructs expressing *PLCG2* variants and the co-factor Rac2^G12V^ ^60^. The assay was conducted using the IP-One Gq kit (Perkin Elmer) according to the protocol. HTRF signal was read by Spark 20M microplate reader (Tecan, 337 nm excitation, 620 nm and 665 nm emission).

## Data sharing

The atomic coordinates and structure factors for the crystal structure of human PLCγ2 (PLCγ2-C) have been deposited in the Protein Data Bank (code 8T7C, http://wwpdb.org/). Structure coordinates for the cryo-EM structures of full-length PLCγ2 (PLCγ2-F), PLCγ2 containing visible H domain (PLCγ2-H), and PLCγ2 in complex with phosphorylated FGFR1K (PLCγ2/pFGFR1K) have been deposited in the Protein Data Bank under entry code 8JQG, 8JQH, 8JQI, respectively. The corresponding cryo-EM maps are deposited in the Electron Microscopy Data Bank under the accession numbers EMD-36571, EMD-36572, and EMD-36573.

## Contributions

Conceptualization YC, ML; investigation YS, AMPM, AM, HC, KT, XZ, JS, NZ, LH, DQ, KS, CX; analysis YS, ML, YC; data curation YS, ML, YC; visualization YS, AM, KT, LH, YC; discussion YS, AMPM, AM, HC, JW, SBP, ML, YC; manuscript preparation YS, ML, YC; supervision YC, ML, JW

## Acknowledgement

We gratefully acknowledge Dr. John Sondek, Dr. Nicole Hajicek, and Dr. Stuart Endo-Streeter for the helpful discussion of PLCγ structure features and FGFR regulation, and advice of FGFR construct design. We gratefully acknowledge Dr. Benjamin Cravatt and Dr. Hui Jing for the helpful discussion and suggestions for the IP One assay. We gratefully acknowledge Dr. Qisheng Zhang for the helpful suggestion of phospholipase activity assay using XY-69.

## Supplementary figure legends

**Supplementary Figure 1.**
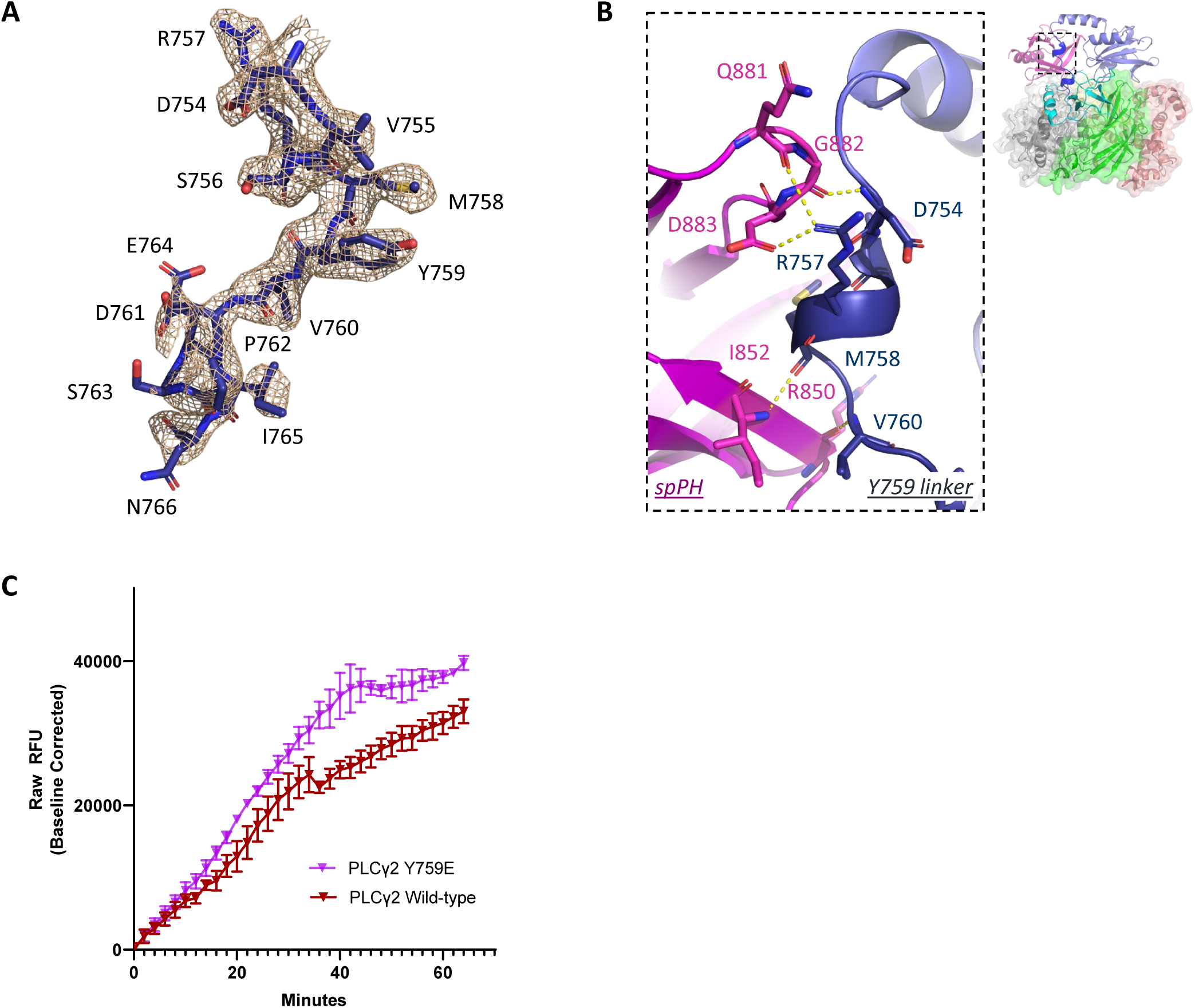
Structure of cSH2-SH3 domain linker containing Tyr759 (Y759 linker, amino acid 754-766) in human PLCγ2. **A.** Electron density and atomic model of Y759 linker. Electron density is rendered at σ = 1.0 and illustrated as wheat-colored mesh. The atomic model is shown in stick representation and the backbone is colored in dark blue. **B.** Polar interactions between cSH2-SH3 linker and spPH domain. Domains are colored as in Figure 1A. Residues contributing to the interactions are shown in stick representation. Interactions are denoted by dashed lines in yellow. Region of representation is illustrated by the dashed box on the complete structure on right. **C.** Biochemical examination of wild-type and Y759E PLCγ2 lipase activity by using recombinant PLCγ2 proteins and a fluorescent substrate mimetic XY-69.

**Supplementary Figure 2.**
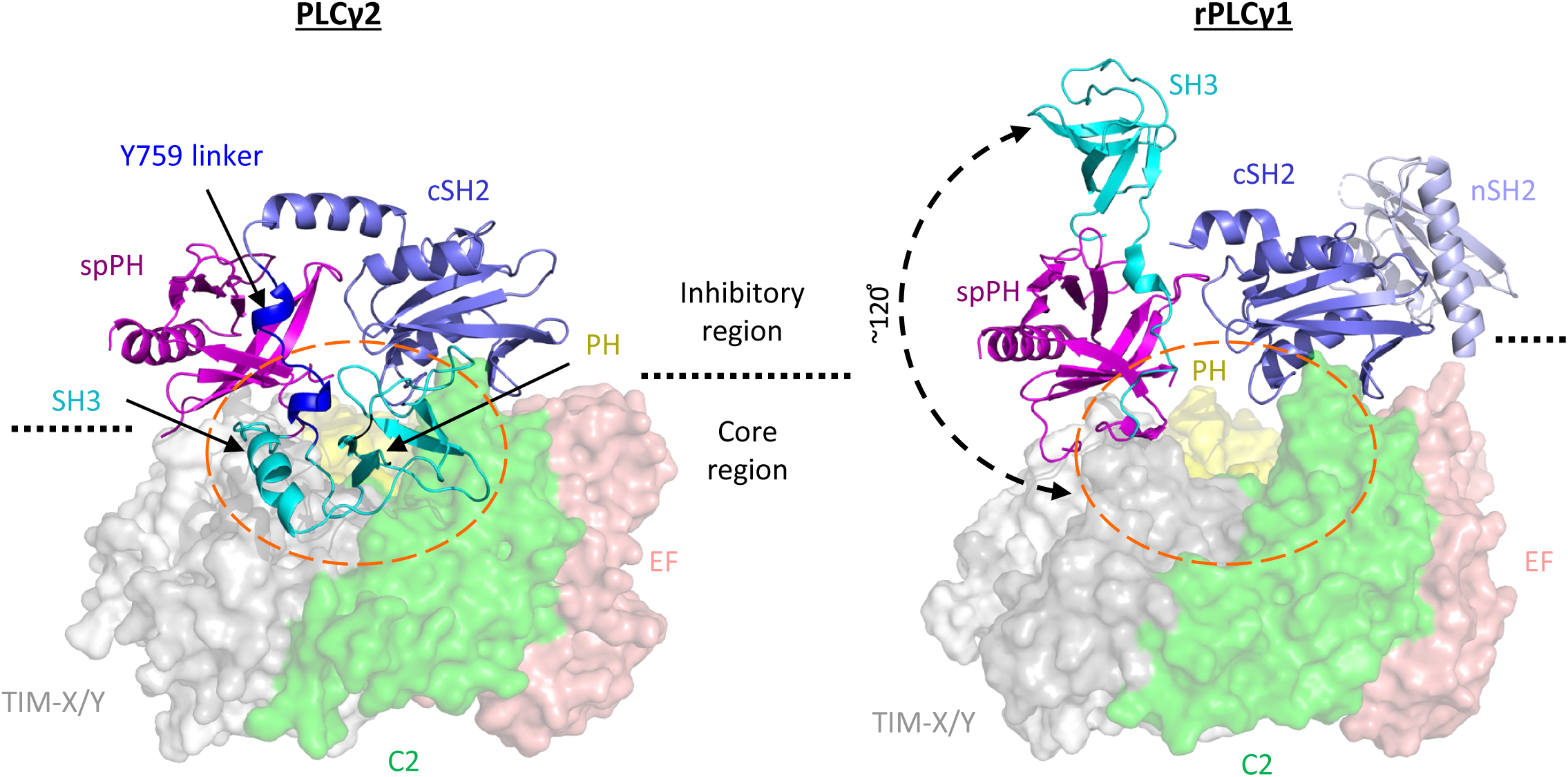
Comparison of the crystal structures of human PLCγ2 and rat PLCγ1 (PDB: 6PBC). Dashed circle in orange highlighted the area occupied by the SH3 domain in PLCγ2 structure. Position difference between the two SH3 domain conformation is demonstrated by dashed arrow in black. Domains are labelled and colored as in Figure 1A. Domains in the inhibitory region are shown in cartoon representation, and core region domains are shown in surface representation.

**Supplementary Figure 3.**
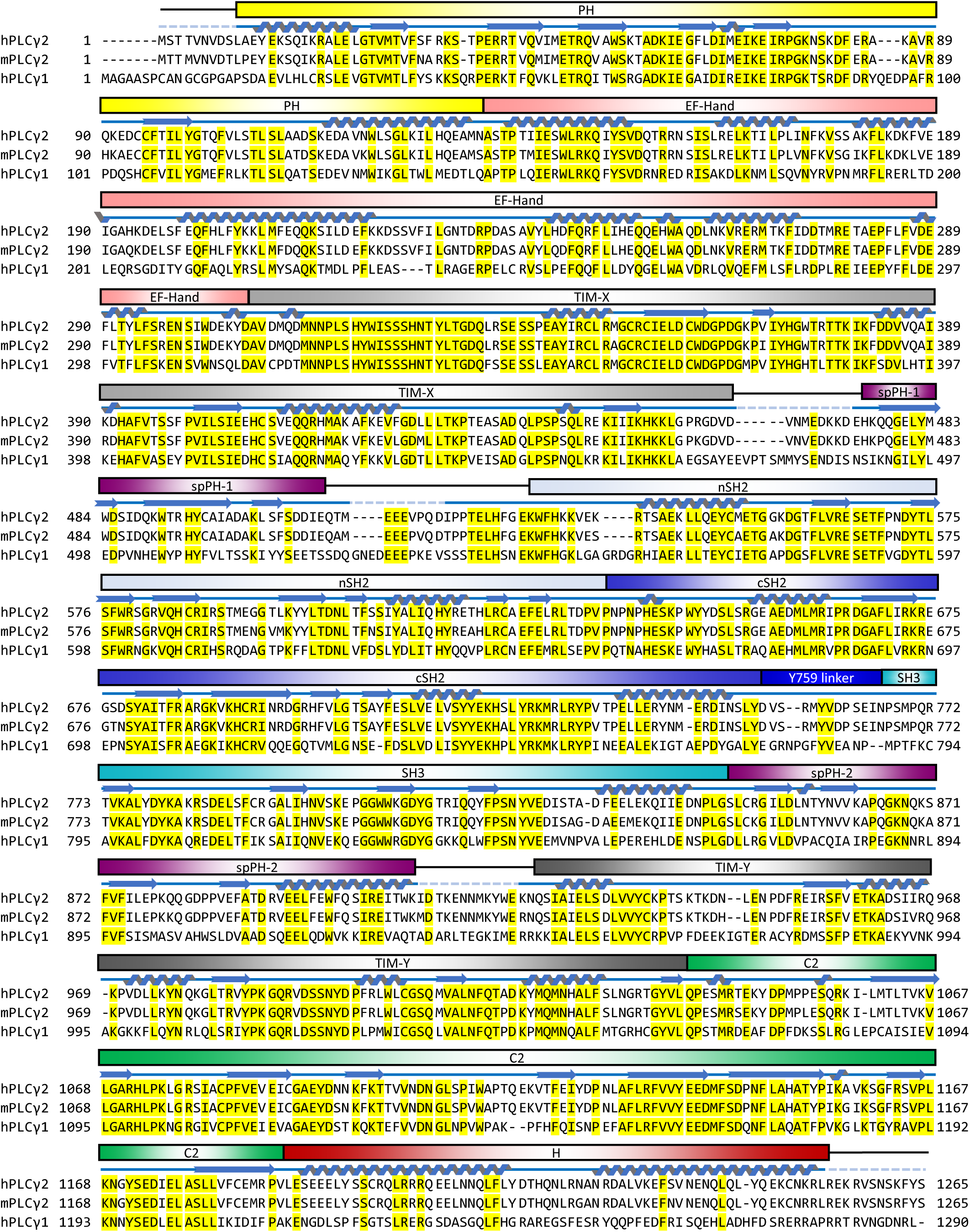
Sequence alignment of human PLCγ2 (hPLCγ2, Uniprot Entry P16885), mouse PLCγ2 (mhPLCγ2, UniProt Entry Q8CIH5), and human PLCγ1 (hPLCγ1, UniProt Entry P19174). Alignment was performed using the Clustal Omega program available in Uniprot. Secondary structures are shown in cartoon representations in blue and domain architectures are annotated as in Figure 1A, both according to the human PLCγ2 structures described in this manuscript. Conserved residues across three sequences are highlighted by yellow background.

**Supplementary Figure 4.**
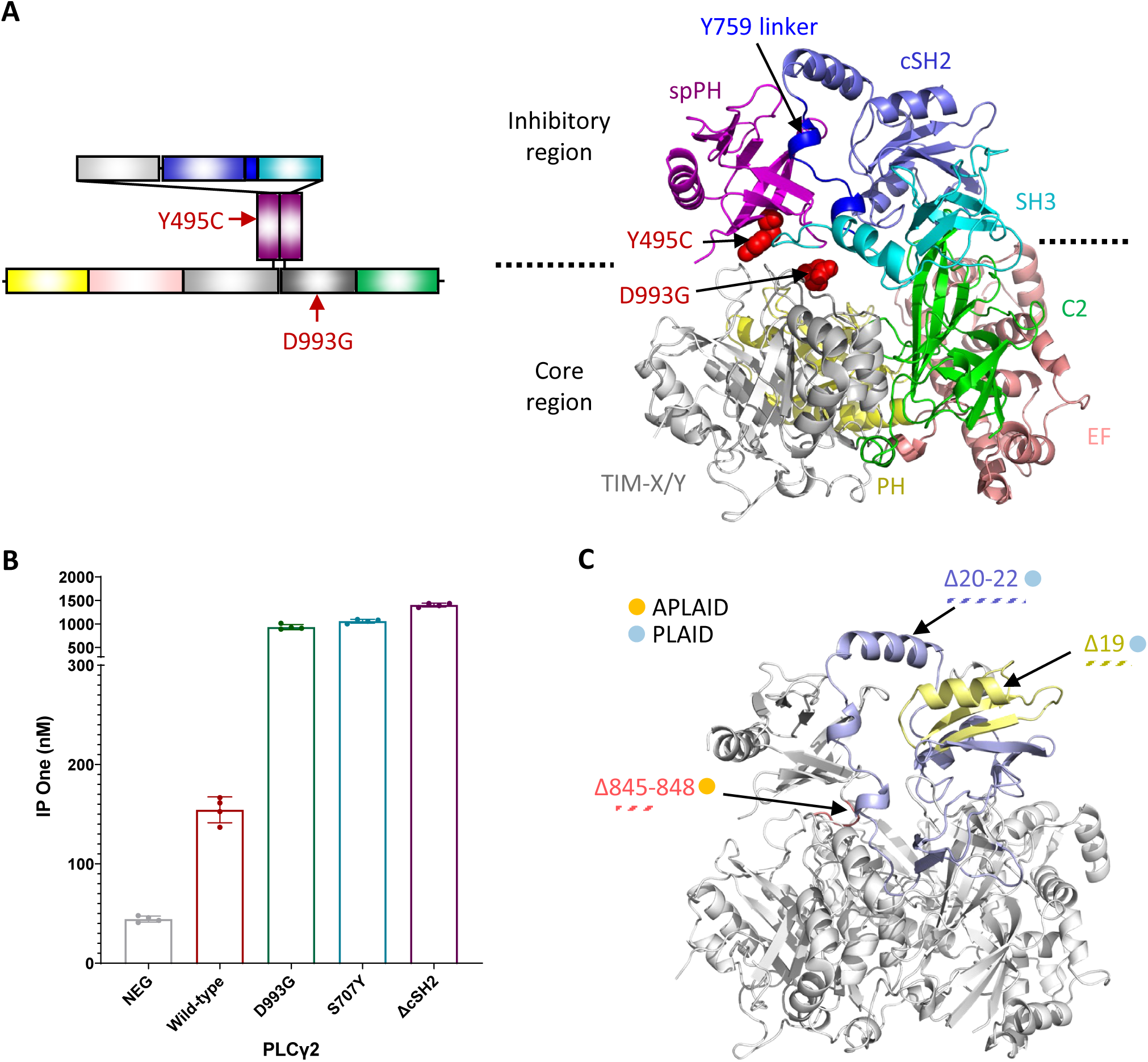
Mapping and activity of PLCγ2 mutations. **A.** Mapping of PLCγ2 mutations from autoinflammatory disease mouse models onto the crystal structure of PLCγ2. Positions of mutations D993G (*Ali5* mice) and Y495C (*Ali14* mice) are denoted by red arrows in the domain illustration cartoon (left) and shown in sphere presentation in red on the crystal structure (right). Domains are colored as in Figure 1A. **B.** Phospholipase activity of PLCγ2 mutations measured in IP One assay. **C.** Mapping of immunodeficiency-associated PLCγ2 deletions onto the crystal structure of PLCγ2. Residues corresponding to the deleted sequences is highlighted by color – yellow: Δ19; blue: Δ20-22; rose: Δ845-848. Association with APLAID/PLAID is illustrated by filled circles as in Figure 2A.

**Supplementary Figure 5.**
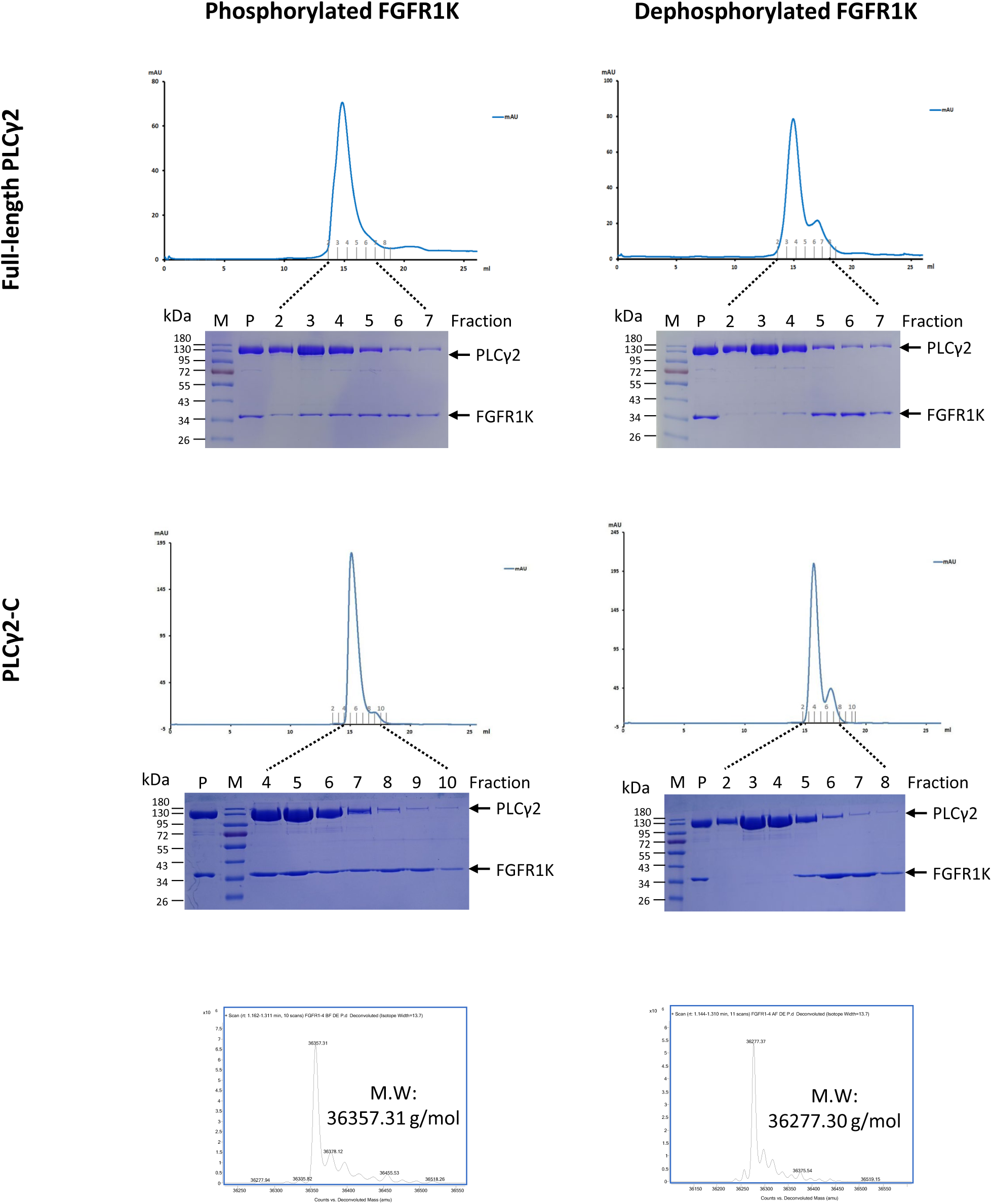
Phosphorylated FGFR1K enhances PLCγ2 association, demonstrated by size-exclusion chromatography. Top: full-length PLCγ2 incubated with phosphorylated (left) and dephosphorylated (right) FGFR1K, respectively. Both size-exclusion chromatography (SEC) elution profiles and corresponding SDS-PAGE examination are shown. Middle: PLCγ2-C incubated with phosphorylated (left) and dephosphorylated (right) FGFR1K, respectively. Bottom: LC-MS analysis of phosphorylated and dephosphorylated FGFR1K, respectively. M: molecular weight marker; P: pre-SEC mixture.

**Supplementary Figure 6.**
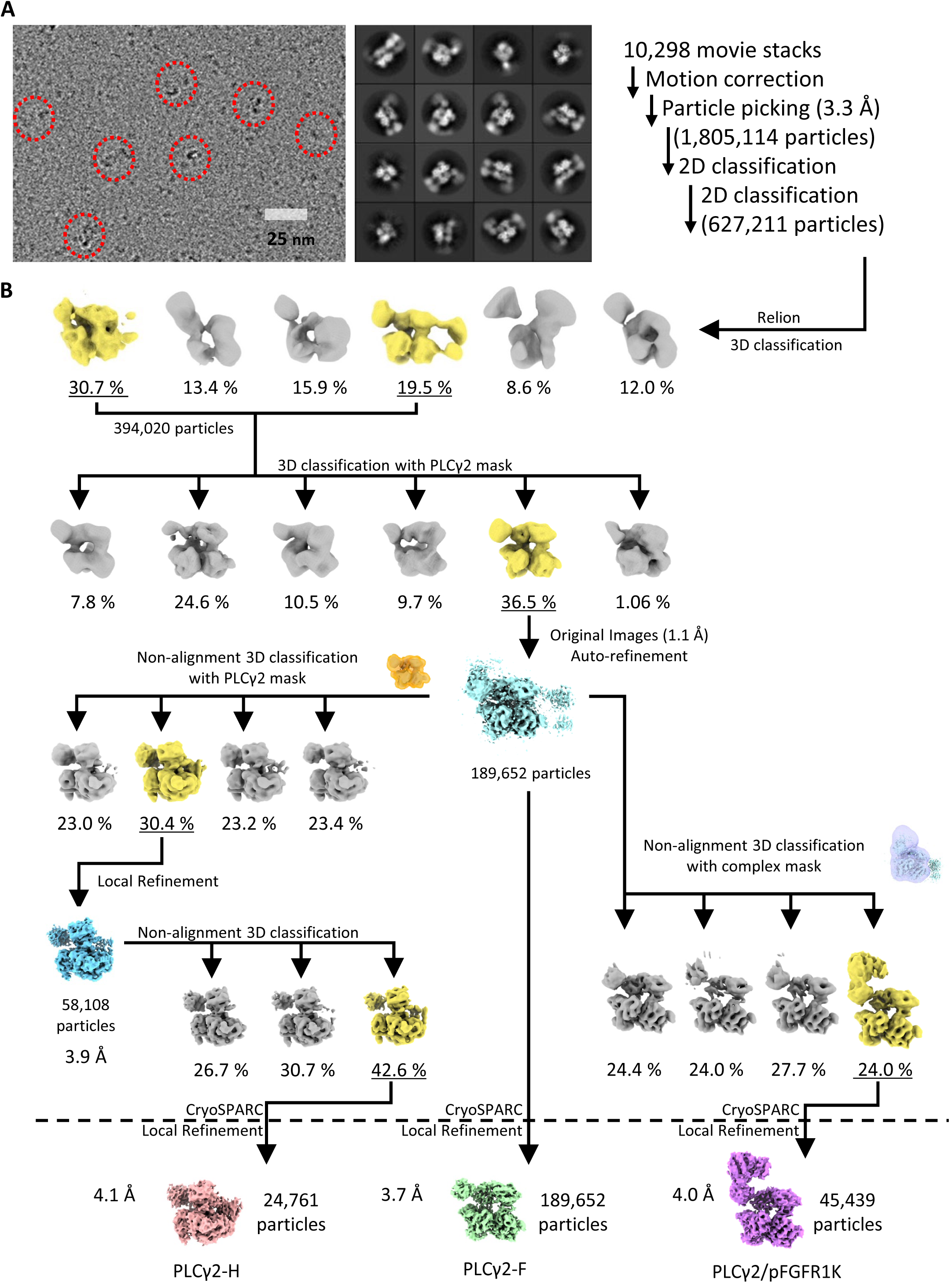
Cryo-EM data processing of full-length PLCγ2 incubated with phosphorylated FGFR1K. **A.** Representative micrograph (left), selected 2D classes (middle), and processing strategy (right) are shown; **B.** Data processing workflow for structure determination of PLCγ2-F, PLCγ2-H, and PLCγ2/pFGFR1K complex.

**Supplementary Figure 7.**
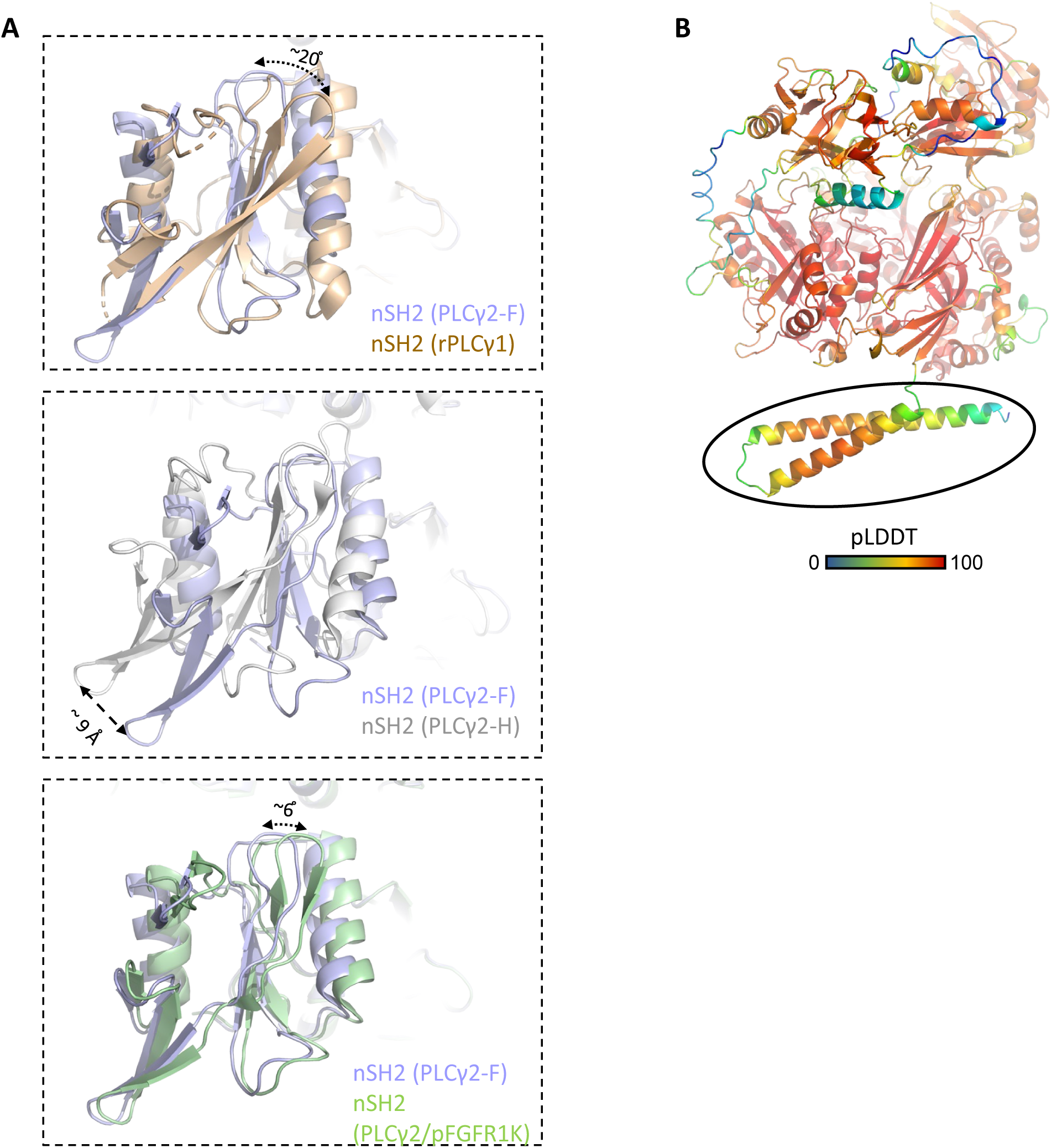
Conformational flexibility of nSH2 domain and AlphaFold predicted model of PLCγ2. **A.** nSH2 domain exhibited conformational flexibility when overlaying PLCγ2 cryo-EM structures, PLCγ2-F, PLCγ2-H, PLCγ2/pFGFR1K, and rat PLCγ1 structure (PDB: 6PBC). **B.** Predicted structure of human PLCγ2 by AlphaFold (AF-P16885-F1) colored by pLDDT value. Predicted helical structures of the C-terminal region is highlighted in the black circle.

## Supplementary tables

**Table S1.**
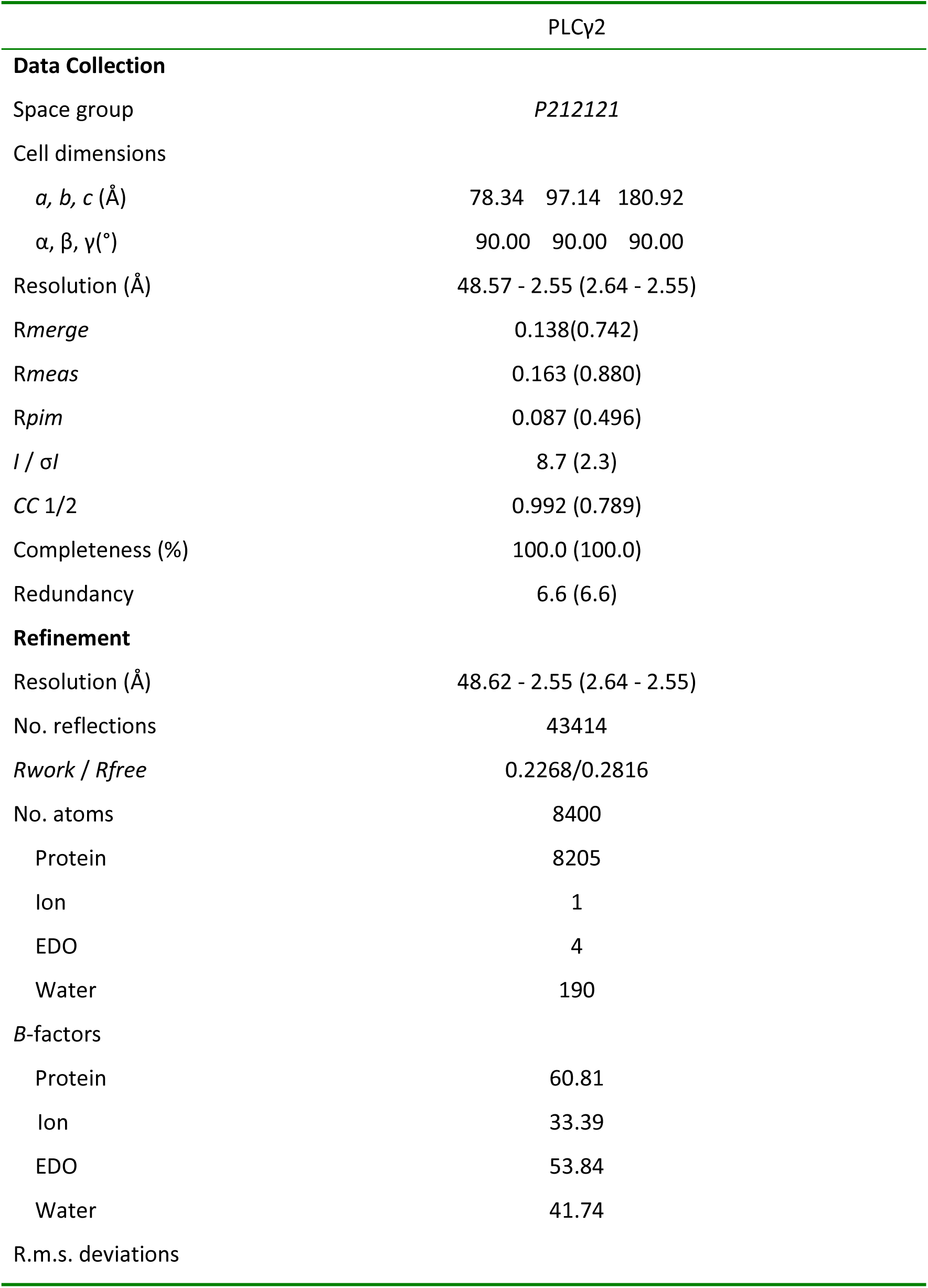
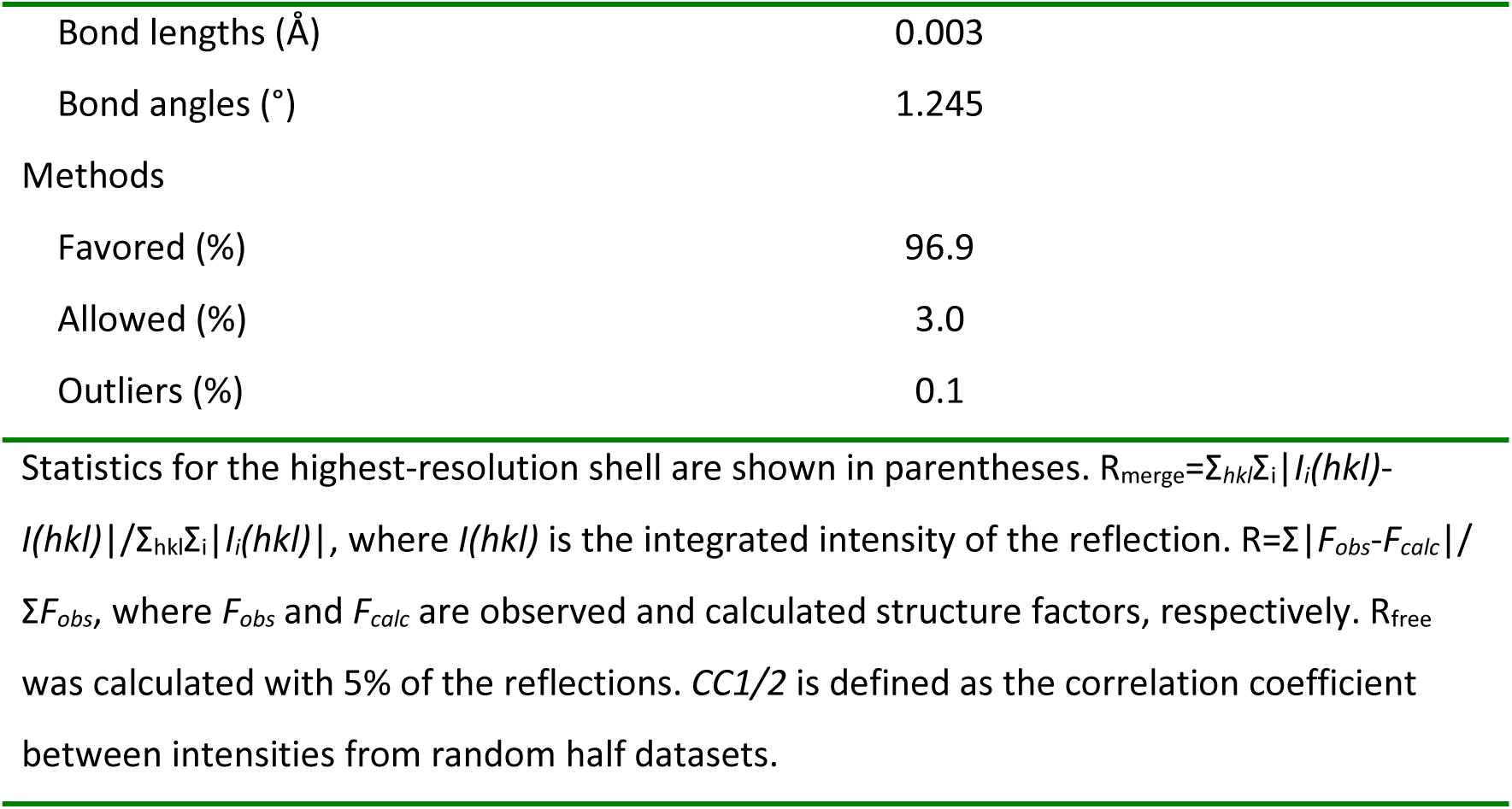
Data collection and refinement statistics.

|  | PLCγ2 |
| --- | --- |
| **Data Collection** |  |
| Space group | *P212121* |
| Cell dimensions |  |
| *a*, *b*, *c* (Å) | 78.34 97.14 180.92 |
| α, β, γ(°) | 90.00 90.00 90.00 |
| Resolution (Å) | 48.57 - 2.55 (2.64 - 2.55) |
| *R*<sub>merge</sub> | 0.138(0.742) |
| *R*<sub>meas</sub> | 0.163 (0.880) |
| *R*<sub>pim</sub> | 0.087 (0.496) |
| *I* / σ*I* | 8.7 (2.3) |
| CC 1/2 | 0.992 (0.789) |
| Completeness (%) | 100.0 (100.0) |
| Redundancy | 6.6 (6.6) |
| **Refinement** |  |
| Resolution (Å) | 48.62 - 2.55 (2.64 - 2.55) |
| No. reflections | 43414 |
| *R*<sub>work</sub> / *R*<sub>free</sub> | 0.2268/0.2816 |
| No. atoms | 8400 |
| Protein | 8205 |
| Ion | 1 |
| EDO | 4 |
| Water | 190 |
| *B*-factors |  |
| Protein | 60.81 |
| Ion | 33.39 |
| EDO | 53.84 |
| Water | 41.74 |
| R.m.s. deviations |  |
| Bond lengths (Å) | 0.003 |
| Bond angles (°) | 1.245 |
| Methods |  |
| Favored (%) | 96.9 |
| Allowed (%) | 3.0 |
| Outliers (%) | 0.1 |
| Statistics for the highest-resolution shell are shown in parentheses. $R_{\text{merge}} = \frac{\sum_{hkl} \sum_i \vert I_i(hkl) - \langle I(hkl) \rangle\vert }{\sum_{hkl} \sum_i I_i(hkl)}$, where $I(hkl)$ is the integrated intensity of the reflection. $R = \frac{\sum \vert F_{\text{obs}} - F_{\text{calc}}\vert }{\sum F_{\text{obs}}}$, where $F_{\text{obs}}$ and $F_{\text{calc}}$ are observed and calculated structure factors, respectively. $R_{\text{free}}$ was calculated with 5% of the reflections. $CC1/2$ is defined as the correlation coefficient between intensities from random half datasets. |  |

**Table S2.**
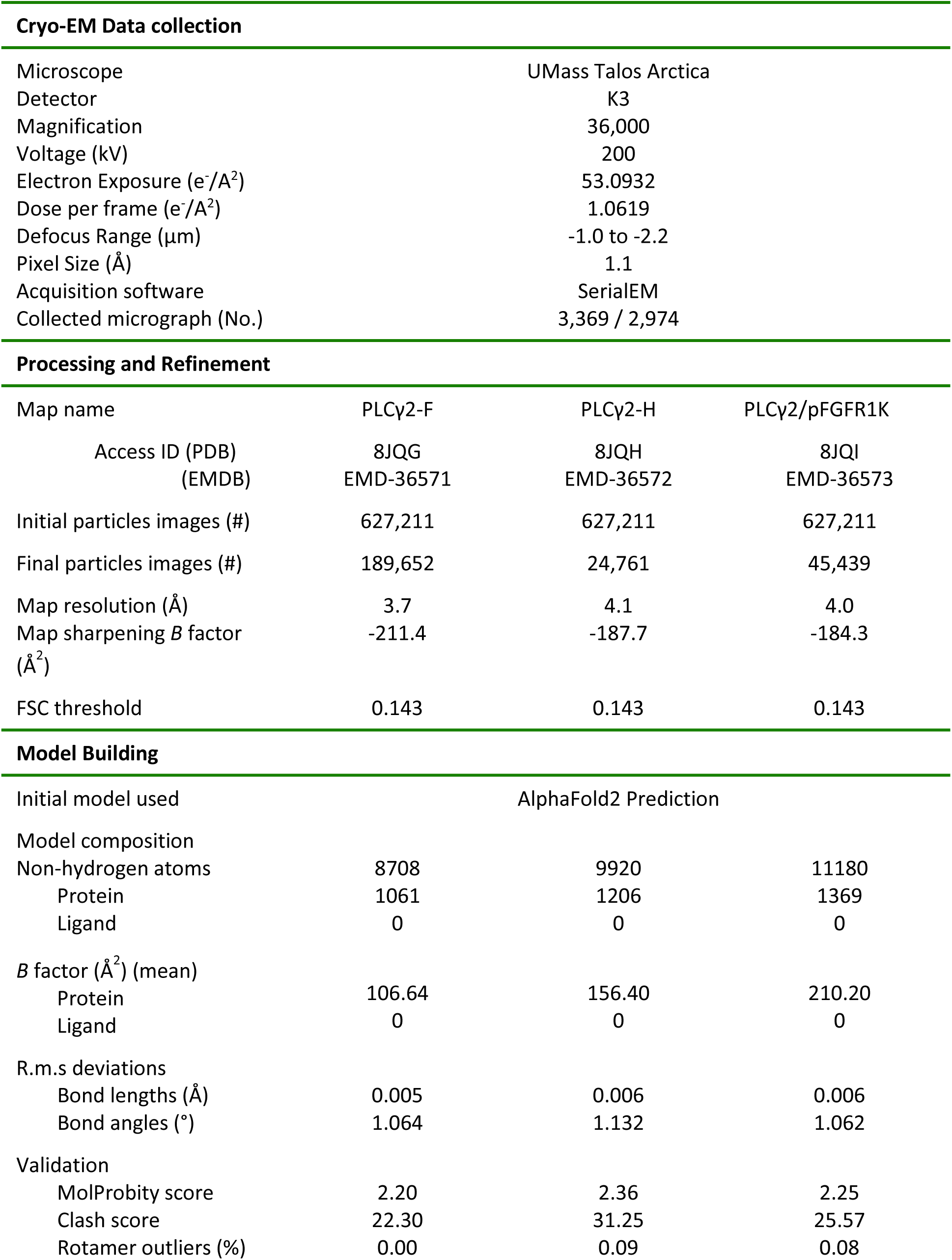
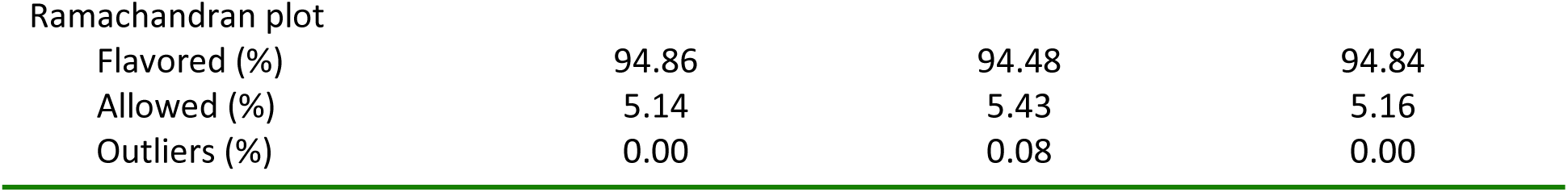
Cryo-EM Data collection, processing, refinement and building statistics.

| Cryo-EM Data collection |  |  |  |
| --- | --- | --- | --- |
| Microscope | UMass Talos Arctica |  |  |
| Detector | K3 |  |  |
| Magnification | 36,000 |  |  |
| Voltage (kV) | 200 |  |  |
| Electron Exposure (e<sup>-</sup>/Å<sup>2</sup>) | 53.0932 |  |  |
| Dose per frame (e<sup>-</sup>/Å<sup>2</sup>) | 1.0619 |  |  |
| Defocus Range (μm) | -1.0 to -2.2 |  |  |
| Pixel Size (Å) | 1.1 |  |  |
| Acquisition software | SerialEM |  |  |
| Collected micrograph (No.) | 3,369 / 2,974 |  |  |
| Processing and Refinement |  |  |  |
| Map name | PLCy2-F | PLCy2-H | PLCy2/pFGFR1K |
| Access ID (PDB) | 8JQG | 8JQH | 8JQI |
| (EMDB) | EMD-36571 | EMD-36572 | EMD-36573 |
| Initial particles images (#) | 627,211 | 627,211 | 627,211 |
| Final particles images (#) | 189,652 | 24,761 | 45,439 |
| Map resolution (Å) | 3.7 | 4.1 | 4.0 |
| Map sharpening *B* factor (Å<sup>2</sup>) | -211.4 | -187.7 | -184.3 |
| FSC threshold | 0.143 | 0.143 | 0.143 |
| Model Building |  |  |  |
| Initial model used | AlphaFold2 Prediction |  |  |
| Model composition |  |  |  |
| Non-hydrogen atoms | 8708 | 9920 | 11180 |
| Protein | 1061 | 1206 | 1369 |
| Ligand | 0 | 0 | 0 |
| *B* factor (Å<sup>2</sup>) (mean) |  |  |  |
| Protein | 106.64 | 156.40 | 210.20 |
| Ligand | 0 | 0 | 0 |
| R.m.s deviations |  |  |  |
| Bond lengths (Å) | 0.005 | 0.006 | 0.006 |
| Bond angles (°) | 1.064 | 1.132 | 1.062 |
| Validation |  |  |  |
| MolProbity score | 2.20 | 2.36 | 2.25 |
| Clash score | 22.30 | 31.25 | 25.57 |
| Rotamer outliers (%) | 0.00 | 0.09 | 0.08 |
| Ramachandran plot |  |  |  |
| Flavored (%) | 94.86 | 94.48 | 94.84 |
| Allowed (%) | 5.14 | 5.43 | 5.16 |
| Outliers (%) | 0.00 | 0.08 | 0.00 |

